# Prediction of Antibody Non-Specificity using Protein Language Models and Biophysical Parameters

**DOI:** 10.1101/2025.04.28.650927

**Authors:** Laila I. Sakhnini, Ludovica Beltrame, Simone Fulle, Pietro Sormanni, Anette Henriksen, Nikolai Lorenzen, Michele Vendruscolo, Daniele Granata

**Affiliations:** Therapeutics Discovery, Novo Nordisk A/S, Copenhagen, Denmark; Digital Chemistry and Design, Novo Nordisk A/S, Copenhagen, Denmark; Centre for Misfolding Diseases, Department of Chemistry, University of Cambridge, UK

**Keywords:** Therapeutic antibodies, non-specificity, protein-language models, machine learning, isoelectric point

## Abstract

The development of therapeutic antibodies requires optimizing target binding affinity and pharmacodynamics, while ensuring high developability potential, including minimizing non-specific binding. In this study, we address this problem by predicting antibody non-specificity by two complementary approaches: (i) antibody sequence embeddings by protein language models (PLMs), and (ii) a comprehensive set of sequence-based biophysical descriptors. These models were trained on human and mouse antibody data from Boughter *et al*. (2020) and tested on three public datasets: Jain *et al*. (2017), Shehata *et al*. (2019) and Harvey *et al*. (2022). We show that non-specificity is best predicted from the heavy variable domain and heavy-chain complementary variable regions (CDRs). The top performing PLM, a heavy variable domain-based ESM 1v LogisticReg model, resulted in 10-fold cross-validation accuracy of up to 71%. Our biophysical descriptor-based analysis identified the isoelectric point as a key driver of non-specificity. Our findings underscore the importance of biophysical properties in predicting antibody non-specificity and highlight the potential of protein language models for the development of antibody-based therapeutics. To illustrate the use of our approach in the development of lead candidates with high developability potential, we show that it can be extended to therapeutic antibodies and nanobodies.

## 1. Introduction

Monoclonal antibodies (mAbs) continue to be one of the leading drug modalities in the pharmaceutical industry, with more than 100 unique mAbs approved by the FDA since 2021^1^ and global sales forecasted to 300 billion US dollars by 2025^2,3^. The success of mAbs for therapeutic application is the result of advances in *in vivo* and *in vitro* discovery platforms, which have enabled fast generation of high-affinity binders towards a highly diverse set of targets^4,5^. Recently, *de novo* design has gained increasing interest in the field as a third-generation discovery approach with the potential of significantly accelerating drug discovery and development timelines^6,7^. Moreover, with the advances in mAb engineering, there has also been an increased interest in the development of mAbs with ultra-high target affinity (pM to fM)^8^, context-dependent target binding (e.g. pH-dependent^9,10^ or ligand induced target binding^11^), and various multi-specific functionalities^12^. To reach optimal binding affinity and potency, mAb hits identified during discovery are often subjected to comprehensive screening campaigns using display platforms (libraries of 10^3^ to 10^10^ variants) of and recombinant well-plate variant generation workflows (libraries of 10^2^ to 10^4^ variants)^13^.

When selecting an antibody lead candidate for development towards clinical testing, optimal target binding affinity and/or pharmacodynamics are key selection parameters. In addition, in the last decade there has been an increased focus on the importance of progressing antibodies to clinical stage which also possess a good developability potential. Antibody developability requires the intersection of multiple disciplines, in which diverse parameters such as expression levels, immunogenicity, processability, and formulation feasibility are addressed, to ensure optimal potential for successful clinical development of lead candidates. Non-specific binding, i.e. weak non-covalent interactions with off-target molecules or interfaces, has emerged as one of the key developability parameters to increase the chance for clinical success^14,15^. Specifically, several studies reported that high tendency for non-specific binding can translate into faster *in vivo* clearance, thereby compromising pharmacokinetics^16-22^. Furthermore, there is an inherent risk that non-specific interactions can translate into undesirable side-effects^23^. Non-specific binding is not a rare phenomenon, as recent reports suggest the presence of a trade-off between affinity and specificity. Thus, optimization of affinity and potency comes with an inherent risk of compromising target specificity^24-27^.

Given the high level of interest in measuring non-specific interactions, there are several *in vitro* screening assays available for this purpose. A commonly used assay is the Enzyme-Linked Immunosorbent Assay (ELISA) with a panel of common antigens, typically insulin, DNA, albumin, cardiolipin and lipopolysaccharide (LPS)^15,27^. Initially, these biomolecules were studied as model antigens for autoimmune responses and diseases. For example, insulin is a self-antigen for autoantibodies associated with type 1 diabetes^28,29^. In addition, DNA, albumin and cardiolipin are self-antigens for autoantibodies associated with several diseases such as systemic lupus erythematosus^30,31^ and anti-phospholipid syndrome^32^, and LPS is an antigen for immune responses to bacterial infections^33^. Moreover, ELISA is widely used in immunology, where non-specific antibodies are often referred to as poly-reactive antibodies. Such antibodies are characterised as having low-affinity binding to multiple distinct antigens, including self-antigens, and they have been widely studied for targets such as HIV and Influenza viruses, as they can be broadly neutralizing^34,35^. While this feature is beneficial for immunity to infectious diseases, to potentially confer broad protection against viruses, it is a highly undesirable feature for therapeutic mAbs. Other common non-specificity assays include baculovirus particle (BVP) ELISA^36^, poly-specific reagent (PSR)^37^, and cross-interaction chromatography with ligands such as heparin^18^ and human IgG from serum^17^.

In addition to these *in vitro* assays, *in silico* methods have been gaining interest, and several tools have been reported recently for prediction of non-specific interactions^38-41^. The development and implementation of predictive computational methods for the prediction and re-design of monoclonal antibody non-specificity at an early stage is of great interest, as it facilitates the generation of safe and efficacious lead candidates with high developability potential. Several studies reported on the identification of non-specificity by *in silico* approaches. Short, linear sequence motifs (e.g. GG, RR, VG, VV, YY, WW and WxW, where x can be any amino acid) enriched in non-specific antibodies, as reported by Kelly *et al*.^25^, have been utilized to create synthetic antibody libraries free from such motifs in the CDRs^42^. Moreover, AI/ML models that classifies non-specific antibodies, leveraging experimental data and sequence-based information, have been reported. Boughter *et al*.^38^ developed a classifier to identify non-specific antibodies based on experimental data acquired from ELISA with a panel of common antigens. Harvey *et al*.^39^ developed a one-hot LogisticReg model based on a naïve Nb library assessed by the PSR assay.

In this study, we developed machine learning (ML) models to estimate the non-specificity of antibodies (**Figure 1A**). Commonly used biophysical properties were tested alongside protein language models (PLMs) to embed antibody sequences. PLMs have emerged as powerful tools for extracting informative features from raw protein sequences by leveraging patterns learned from massive sequence databases^43-46^. Among these, Evolutionary Scale Modeling (ESM) models have shown particular promise in capturing structural, functional, and physico-chemical properties (including antibody specificity) without requiring explicit structural data^47-50^. ESM models, such as ESM-1v, encode sequences into high-dimensional embeddings that reflect residue context, conservation, and evolutionary information, which are all factors known to influence antibody behaviour. These features make ESM models well-suited for predicting complex properties like non-specificity, which can arise from subtle sequence-dependent effects not easily captured by traditional descriptors. By applying ESM models to antibody variable regions, we aim to harness its representation power to identify sequence signatures of non-specific binding and improve early-stage developability assessment.

**Figure 1.**
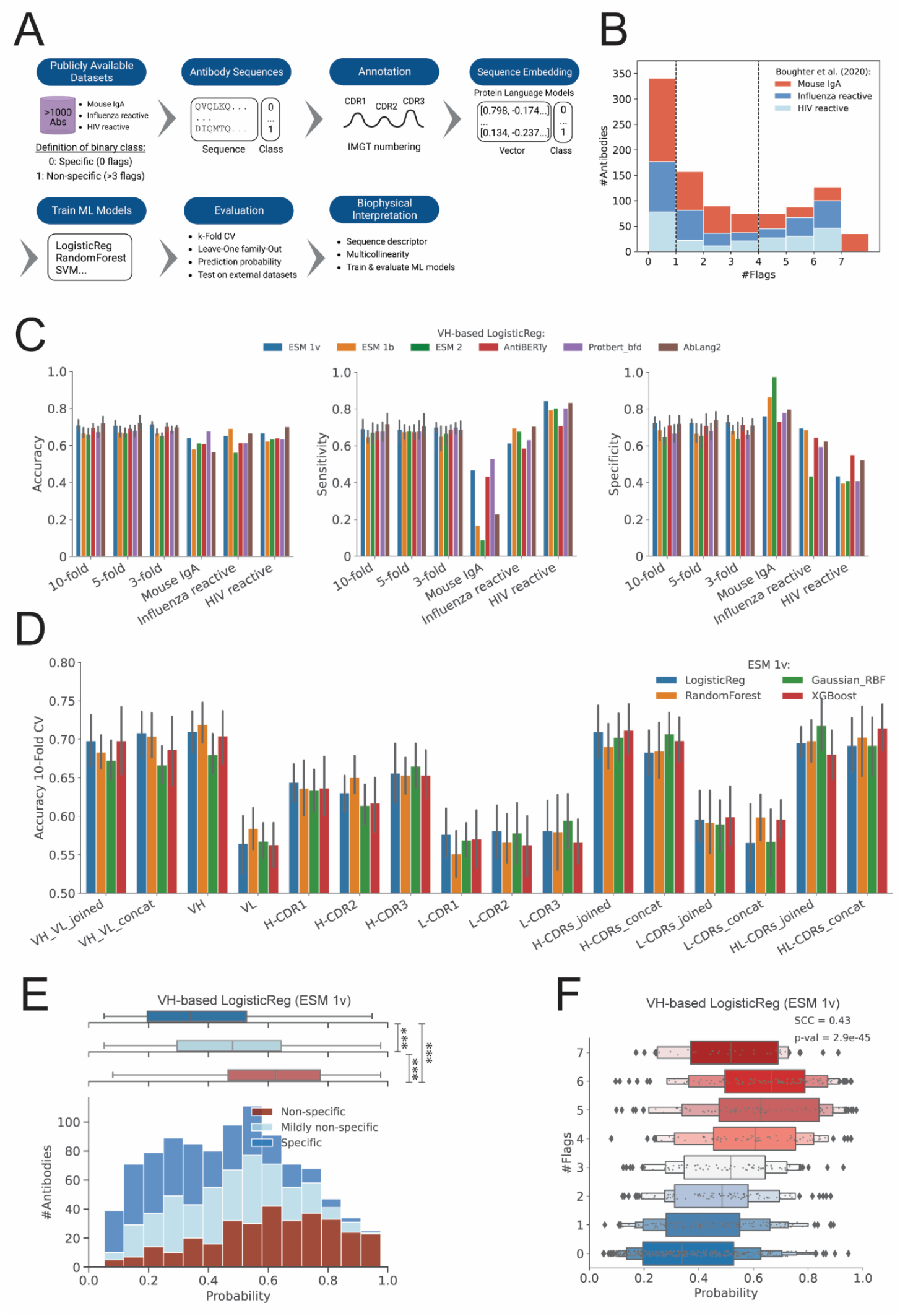
Performance Evaluation of Machine Learning Models for Predicting Antibody Non-Specificity. **(A)** Schematic workflow of the study. Publicly available datasets containing antibody sequences were used. These sequences were annotated andembedded using sequence-based biophysical descriptors and protein language models (PLMs). Different ML models were trained and evaluated using k-fold cross-validation, sensitivity-specificity analysis, and external datasets. **(B)** Histogram showing the distribution of antibody sequences based on the number of flags in the Boughter dataset. Sequences are categorized as Influenza reactive (blue), HIV non-reactive (red), and HIV reactive (light blue). **(C)** Bar plots showing the validation performance (k-Fold CV and Leave-One Family-Out) for a top performing ML algorithm (LogisticReg) and PLMs across various validation schemes (k-fold CV and leave-one-family-out). **(D)** Bar plot of 10-fold CV accuracy for different antibody sequences embedded by top performing language model (ESM 1v mean-mode). **(E)** Histogram of predicted probabilities of antibody non-specificity using the VH-based Logistic Regression (ESM 1v) model for the Jain dataset. Antibodies are classified into non-specific (dark blue), mildly non-specific (light blue), and specific (red) categories. **(F)** Boxplot comparing the predicted non-specificity probabilities for antibodies across different datasets using VH-based Logistic Regression (ESM 1v). The boxplot displays the median, interquartile range, and outliers, with significant differences indicated by SCC and p-values (*** indicate p-value <0.001).

Besides testing which encoding provided the best prediction performance, one aspect of the study was to identify which part of the antibody contributes to non-specificity. Furthermore, to gain biophysical insight, sequence-based biophysical descriptors were analysed to support the predictive models. Our results indicate that the computational models that we considered enable the prediction of non-specific interactions that can be used to guide the design and selection of antibodies with improved specificity and efficacy.

## 2. Results & Discussion

### 2.1 Public antibody data

Four different datasets were retrieved from public sources; (i) a curated dataset of >1000 mouse IgA, influenza-reactive and HIV-reactive antibodies with their respective non-specificity flag from ELISA with a panel of common antigens^38^, (ii) 137 clinical-stage IgG1-formatted antibodies with their respective non-specificity flag from ELISA with a panel of common antigens^15^, (iii) 398 antibodies, originating from naïve, IgG memory and long-lived plasma cells, with their respective poly-specific reagent score^51^, and (iv) 140 000 nanobody (Nb) clones assessed by the PSR assay from a naïve Nb library^39^. These four datasets are referred to as the Boughter, the Jain, the Shehata and the Harvey datasets, respectively.As therapeutic antibodies are engineered to be closely related to human antibodies to avoid immunogenic responses, it is important to exploit human antibody data for development of optimal ML models. As the Boughter dataset partly consists of mouse IgA antibodies, the sequence similarity of these mouse antibodies was compared to the human antibodies to ensure that there are not too large sequence differences within the dataset. The mouse IgA antibodies appear to differ mostly in the H/L-CDRs (**Figure S1B**). Another notable difference within the Boughter dataset is that the mouse IgA antibodies have a slightly shorter H-CDR3 loop relative to the human antibodies (**Figure S1C**).

Ideally, training data sets for classification ML-Models should be balanced when it comes to positive and negative data points. The distribution of non-specificity for the three datasets is visualised in **Figure 1B**. The Boughter and the Jain datasets are relatively balanced in terms of specific (zero flags), mildly non-specific (1-3 flags) and non-specific (>4 flags) antibodies, while the Shehata dataset is unbalanced, with 7 out of 398 antibodies characterised as non-specific only. In this study, the most balanced dataset (i.e. Boughter one) was selected for training of ML models, while the remining three (i.e. Jain, Shehata and Harvey, which consists exclusively of VHH sequences) were used for testing.

### 2.2 Protein language models enable the representation of antibody non-specificity

Following the study original study^38^, the Boughter dataset was first parsed into two groups: specific (0 flags) and non-specific group (>3 flags), leaving out the mildly non-specific antibodies (1-3 flags) (**Figure 1A**). The amino acid sequences of the parsed dataset were then annotated in the CDRs, and various fragments of the antibody sequences were embedded into vectors representing their physico-chemical and structural properties (i.e. ESM 1b, ESM 1v, ESM 2, Protbert bfd, AntiBERTy, and AbLang2). This procedure resulted into the training of 12 different antibody fragment-specific binary classification models were trained (see **Table 4**). The performance of all the generated models from 10-fold cross-validation (CV) can be seen in **Figure S2-S8**. Overall, all of the protein language models (PLMs) performed well with 66-71% 10-fold CV accuracy, including the antibody-specific ones AntiBERTy and AbLang2 **(Figure 1C**). These deep learning models were trained on large datasets of protein sequences in the million-to-billion range, encoding protein sequences into vectors for representation of their physiological properties, remote homology, and secondary/tertiary structure. PLMs were originally developed for the prediction of protein contacts and structure^52,53^.

Going forward, the Evolutionary Scale Modelling (ESM) 1v was selected as the embedder of choice for this study.

### 2.3 The highest PLM-based predictability is achieved by encoding the VH domain

An overview of the different antibody fragment-specific models based on the ESM 1v embedder is shown in **Figure 1D**. Highest predictability (71% 10-fold CV accuracy of non-specificity) was obtained for the models trained on the VH and H-CDRs sequences. These results suggest that the non-specificity primarily originates from the VH domain, with main contributions from the H-CDR loops. When looking at the models based on the individual H-CDR loops, the order of low-to-high predictability of non-specificity follows H-CDR2, H-CDR1 and H-CDR3 (**Figure 1D**). H-CDR3 has the highest predictability of non-specificity among all the H-CDR loops. The importance of the H-CDR3 loop for non-specificity is in agreement with Guthmiller *et al*.^35^, who showed by using MD simulation that the flexibility of the H-CDR3 loop plays an important role in the non-specific behaviour of antibodies.

Accuracy of around 70% was consistently observed across 3, 5 and 10-Fold CV for the top performing models (**Figure 1C**), and similar performance obtained for sensitivity and specificity. Moreover, when looking at the predictability of one antibody family to another, the overall accuracy was consistently above >60% for the Leave-One Family-Out validations. However, when comparing sensitivity and specificity, classifiers trained on human antibodies performed poorly when tested on mouse antibodies. This is not surprising as mouse and human antibodies have notable sequence differences, such as mouse IgA having a shorter H-CDR3 and larger sequence differences in the CDRs relative to human antibodies (**Figure S1B**). Moreover, classifiers trained on mouse IgA and HIV reactive antibodies perform well across all evaluation metrics (accuracy, sensitivity and specificity) when tested on Influenza reactive antibodies, while classifiers trained on mouse IgA and Influenza reactive antibodies seem to be better in predicting non-specific HIV reactive antibodies than specific ones.

### 2.4 Classification probability of non-specificity against non-specificity ELISA flags mimics regression behaviour

A prediction probability of non-specificity was computed in addition to the binary output from the binary classification models. When comparing the prediction probability of non-specificity for all the antibodies from the Boughter dataset (test antibodies sampled from the 10-Fold CV and the mildly non-specific antibodies), three distinct distributions of prediction probabilities for the specific, mildly non-specific and non-specific antibody groups appeared (**Figure 1E**). The premise that non-specificity is not a binary property is exemplified by the overlaps between the distributions. This illustrates that the prediction probability can be used beyond the binary output to assess antibodies of varying degree of non-specificity.

When comparing those to non-specificity ELISA flags, a significant regression-like behaviour (SCC 0.43) was observed for one of the top performing classifiers, ESM 1v mean-mode VH-based LogisticReg model (**Figure 1F)**. The prediction probability for non-specificity followed an uptrend when compared to the non-specificity ELISA flags. An exception to this trend was the antibodies with seven non-specificity ELISA flags, as those were exclusively mouse IgA antibodies (differences discussed in previous section).

### 2.5 The isoelectric point: a key biophysical driver of non-specificity

To gain insight into the biophysical origins of antibody non-specificity, a set of 68 sequence descriptors (**Table S1**) was computed for the parsed Boughter dataset. These descriptors encompass a wide range of biophysical properties derived from the antibodies sequence, including theoretical isoelectric point (pI), secondary structure propensity, and hydrophobicity. To assess the presence of redundancy among the descriptors, we constructed a Spearman’s correlation matrix, which revealed that several descriptors, such as hydrophobicity descriptors, exhibited strong correlation among each other (SCC > 0.5, **Figure S9**), thus indicating redundancy. All the descriptors were ranked according to the absolute logistic regression coefficients (**Table S2**), whereafter top 25 descriptors were selected, and used for training of a VH-based LogisticReg model. The in-depth analysis of the importance of the 25 descriptors is shown in four different plots in **Figure 2A**:

**Figure 2.**
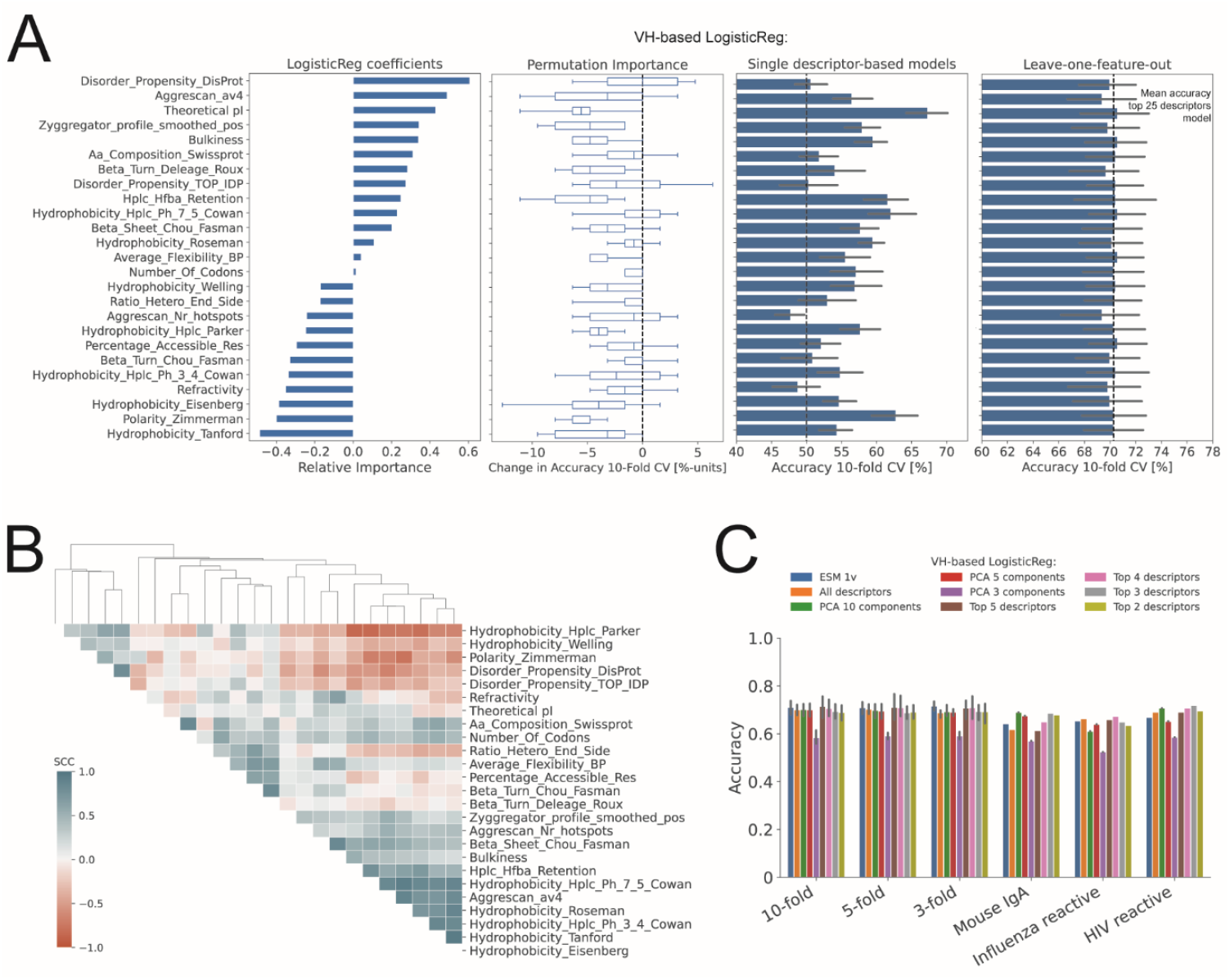
Analysis of Descriptor Importance and Model Performance for VH-Based Logistic Regression. **(A)** Analysis of descriptor importance using various metrics for the VH-based Logistic Regression model; (first panel) Logistic regression coefficients indicating the relative importance of different features, (second panel) permutation importance showing the change in 10-fold CV accuracy when each descriptor is permuted, (third panel) 10-fold CV accuracy of models based on single descriptors, and (fourth panel) 10-fold CV accuracy of leave-one-feature-out models compared to the mean accuracy of model using the top 25 descriptors. **(B)** Heatmap displaying the Spearman’s correlation coefficient (SCC) between the top 25 descriptors selected based on highest Logistic Regression coefficient in model with all descriptors. The dendrogram shows hierarchical clustering of descriptors based on their SCC. **(C)** Bar plot comparing the 10-fold CV accuracy of different models in predicting antibody non-specificity across various validation schemes: k-fold CV and leave-one-family-out validation. Models compared include VH-based sequences embedded by ESM 1v, all descriptors, PCA with 3, 5 and 10 components, and top 2, 3, 4, and 5 descriptors. Sensitivity and specificity bar plots can be found in **Figure S12**.

- The first plot shows the LogisticReg coefficients, indicating the relative importance of each descriptor. Notably, Disorder_Propensity_DisProt, Aggrescan_a4v, and theoretical pI show significant positive coefficients, suggesting that they are strong drivers of non-specificity.
- The second plot displays the permutation importance, highlighting the change in accuracy when each descriptor is permuted. Descriptors like theoretical pI, bulkiness, Hplc_Hfba_retention and Polarity_Zimmerman demonstrate substantial decrease in accuracy upon permutation.
- The third plot illustrates the accuracy of models based on single descriptors to underscore their individual predictive power. Theoretical pI resulted in the highest accuracy compared to the other descriptors, confirming its critical role in the prediction of non-specificity.
- The fourth plot shows the leave-one-feature-out accuracy, revealing how the exclusion of each descriptor affects the overall model performance. Most of the descriptors result in a minimal drop in accuracy, indicating that the model performance remains unaffected when a certain descriptor is left out. This can be explained by that there remains a certain level of redundancy among the 25 descriptors, e.g. theoretical pI appears to be negatively correlated with Polarity_Zimmerman, according to the Spearman’s correlation matrix in **Figure 2B**

To further narrow down the redundancy, the 25 descriptors were tested in all possible combinations of 2, 3, 4 and 5 descriptors for training of new LogisticReg models. The results of the top descriptors from this analysis are shown in

**Table** 1. The results indicate that, among the top 5 descriptors, the theoretical pI appear to be the most important driver for non-specificity. This conclusion is also supported by the frequency of this particular descriptor among the top models (**Figure S10**). Additionally, as a parallel check, we performed Principal Component Analysis (PCA) for dimensionality reduction and feature selection. The primary objective of this study was again to address multicollinearity among the descriptors and to identify the most significant features contributing to the variance in the dataset. We thus evaluated the performance of the LogisticReg models trained on the 3, 5, and 10 principal components identified by PCA. In agreement with our previous findings regarding the theoretical pI, the presence of this descriptor among the selected features significantly influenced the model performance, particularly when it was included in PCA 5 and 10 components (see **Figure 2C**). The importance of theoretical pI was captured at ≥5 components, as seen by the magnitudes of Eigenvalues of theoretical pI for PCA 5 as compared to PCA 1-4 (see **Figure S11**). The isoelectric point is known to influence PKPD behaviour/clearance of antibodies^54,55^.

Altogether, a comparison between the PLM-based and descriptor-based ML models in terms of accuracy of across different validation schemes is shown in **Figure 2C**. The results indicate that the ESM 1v model consistently achieves high accuracy across all validation schemes. The VH-based Logistic Regression model using all descriptors also performs well, though slightly lower than the ESM 1v model. Notably, the PCA-based models show comparable performance, demonstrating the effectiveness of dimensionality reduction in maintaining model accuracy. Interestingly, the VH-based Logistic Regression models using the top descriptors, 2, 3, 4, and 5 combinations (**Table** 1), exhibit robust performance. This finding suggests that a smaller subset of key descriptors can achieve similar predictive power as using the full set of descriptors, highlighting the potential for model simplification without compromising accuracy.

**Table 1.** Top 2, 3, 4 and 5 combined VH-based sequence descriptors.

| Top | Descriptors |
| --- | --- |
| 2 | Theoretical pI, Bulkiness |
| 3 | Theoretical pI, Bulkiness, Aggrescan_Nr_hotspots |
| 4 | Theoretical pI, Average_Flexibility_BP, Beta_Turn_Chou_Fasman, Hplc_Hfba_Retention |
| 5 | Theoretical pI, Bulkiness, Disorder_Propensity_DisProt, Percentage_Accessible_Res, Aggrescan_av4 |

### 2.6 VH-based LogisticReg classification model is applicable to clinical-stage therapeutic antibodies

To show applicability of the non-specificity classification model on therapeutic antibodies, the ESM 1v mean-mode VH-based LogisticReg model was tested on the Jain dataset. As in the Boughter dataset, the Jain dataset was parsed into two groups, specific (0 flags) and non-specific (>3 flags), leaving out the mildly non-specific antibodies (1-3 flags). An accuracy of 69% was obtained for the parsed Jain dataset (see confusion matrix in **Figure S14A**). This value is comparable to the mean accuracy of 71% obtained for the same classifier across 3, 5 and 10-Fold CV for the parsed Boughter dataset. Moreover, as in the case of the Boughter dataset, a similar distribution of prediction probability of non-specificity was obtained for the full Jain dataset, and it appears to mimic regression-like behaviour when compared to the non-specificity ELISA flags, although to a slightly weaker extent (**Figure 3A** and **S13**). The same trend can be observed for the top 5 descriptors model (**Figure 3C**). The overall performance of the classifier on the Jain dataset illustrates that it can be applied to therapeutic antibodies.

**Figure 3.**
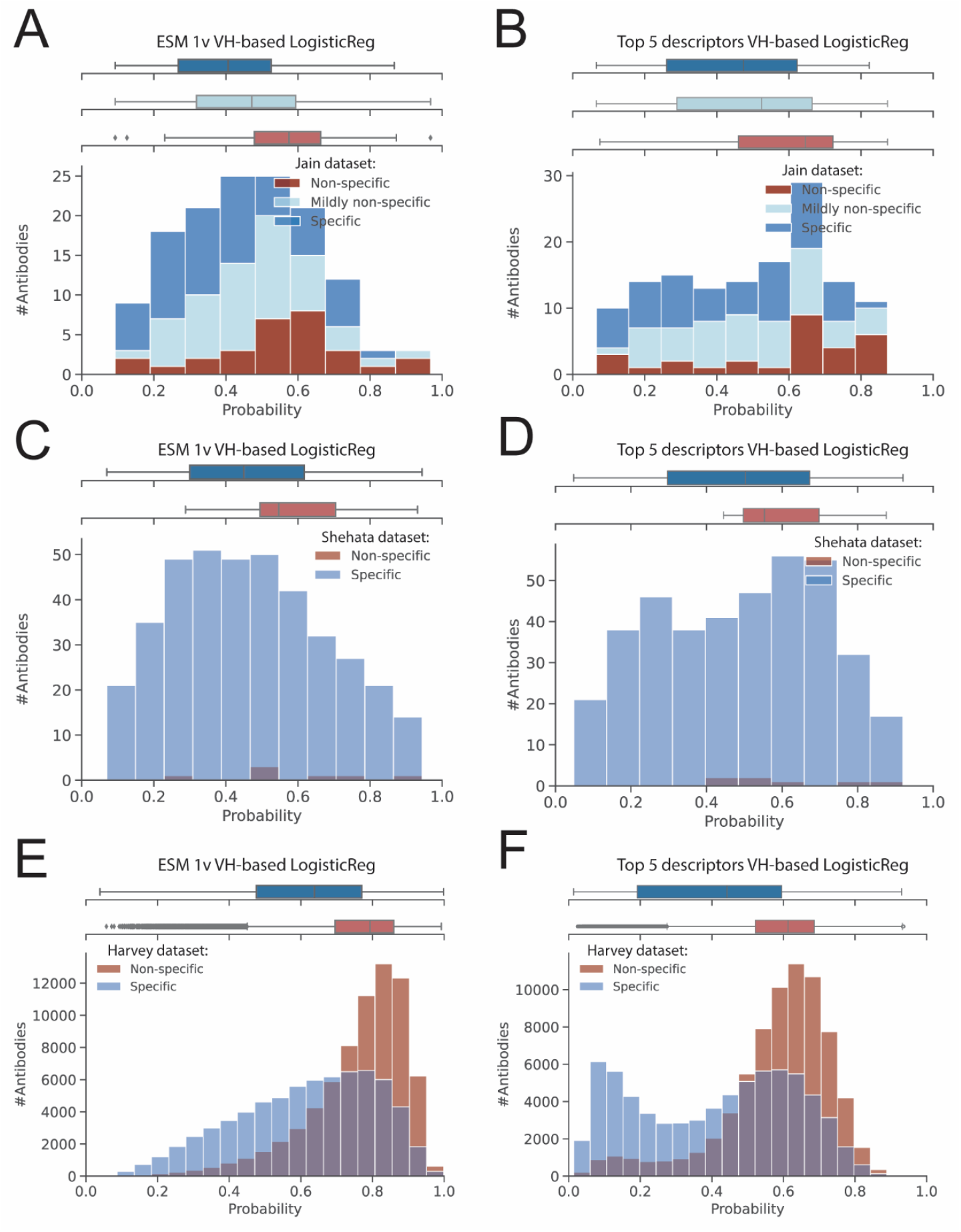
Logistic Regression Models Predicting Antibody Non-Specificity Across Different Datasets. **(A-F)** Distributions of predicted probabilities of antibody non-specificity for three different datasets using two logistic regression models: predictions for the Jain dataset (A,B), for the Shehata dataset (C,D), and for the Harvey dataset (E,F). (A, C, and E) depict results from the ESM 1v VH-based logistic regression model, while (B, D, and F) depict results from the top 5 descriptors VH-based logistic regression model. For each dataset, antibodies are classified into specific, mildly non-specific (only in the Jain dataset), and non-specific categories, represented by different colours.

### 2.7 Antibodies characterised by the PSR assay appear to be on a different non-specificity spectrum than that from the non-specificity ELISA assay

During recent years, alternative assays to ELISA have been developed to meet the demand of high-throughput screening during drug discovery, and such one is the poly-specific reagent (PSR) assay^37^, where antibodies displayed on the surface of yeast cells are counter-selected when non-specifically bound to soluble membrane protein in a flow cytometry-setup. Several studies have been reported using this assay for assessing antibody non-specificity^15,17,51^. Recently, Harvey and co-authors^39^ developed a one-hot LogisticReg model based on >140 000 clones assessed by the PSR assay from a naïve Nb library. They found a significant correlation of the PSR with the gold-standard ELISA based on six Nbs. To find out whether our ESM 1v mean-mode VH-based LogisticReg model can extend its applicability further to the non-specificity scored by the PSR assay, the Shehata dataset and the VH-based Nb dataset by Harvey and co-authors^39^, here referred to as the Harvey dataset, were tested. The classifier did not appear to separate the PSR-scored specific and non-specific antibodies well. All the specific PSR-scored antibodies of the Shehata dataset were distributed along the entire prediction probability scale, while the few non-specific ones were on the probability end towards higher non-specificity (**Figure 3C,D**). A similar forecast was observed for the Harvey dataset; all the specific PSR-scored Nbs resulted in a broad probability distribution, while the non-specific PSR-scored ones resulted in a narrower probability distribution towards higher non-specificity (**Figure 3E,F**). Thus, the classifier appears to be better at predicting non-specific PSR-scored antibodies, than specific PSR-scored antibodies. This result suggests that the spectrum of non-specificity from the PSR assay is different than the one from the non-specificity ELISA assay, of which the classifier is trained on. Thus, a specific antibody classified by the PSR assay may necessarily not translate into a specific antibody classified by the non-specificity ELISA assay. The specific PSR-scored Nbs could partly consist of mildly non-specific clones in addition to specific ones, thus resulting in this broad probability distribution.

An interesting remark can be made about the distributions of predicted probabilities obtained from the two different LogisticReg models tested on the Harvey dataset in **Figure 3E,F**. The ESM 1v VH-based LogisticReg model produces a more uniform distribution of predicted probabilities across the dataset, while the top 5 descriptors VH-based LogisticReg model exhibits a clear biphasic distribution. It is no surprise that this bimodal pattern closely resembles the distribution of pI (**Figures S15-S18**), as this descriptor is the main driver of non-specificity in the top 2, 3, 4 and 5 descriptors LogisticReg models (**Table** 1 and **Figure S10**). Nonetheless, the distribution of non-specific antibodies in the Harvey dataset appears to exclusively be of high pI (>8) according to **Figure S18A**. The distributions of the other descriptors do not appear to differ significantly between specific and non-specific antibodies see (**Figures S15-S18**). The distinct separation suggests that the pI plays a crucial role in differentiating between specific and non-specific antibodies.

### 2.8 VH-based LogisticReg models performs on par or better than existing predictors

The ESM 1v VH-based LogisticReg model can be compared to two existing predictors in the literature - the predictors reported in the Boughter *et al*. study^38^, and the Harvey *et al*. study^39^. Boughter and colleagues stated that while no notable difference could be observed between specific and non-specific antibodies in both gene usage level and amino acid-usage level in CDRs, the positional context of biophysical properties can show the differences. They showed to be successful in developing a binary classifier based on a position-sensitive biophysical matrix with accuracy up to 75%. Their reported performance is on par with the achieved performance of the PLM-based classifier (ESM 1v VH-based LogisticReg), which was trained on the same data (parsed Boughter dataset).

Furthermore, Harvey and colleagues developed a one-hot LogisticReg model based on >140 000 clones assessed by the PSR assay from a naïve Nb library with an accuracy >80%. Using their published web-based predictor^56^, we tested its performance on the Boughter, Jain, and Shehata datasets. The results show that the Harvey predictor does not separate well the different antibody groups in the Boughter dataset (**Figure S19A**,**B**), with overlapping distributions of prediction scores. Similarly, the Jain and the Shehata datasets demonstrate significant overlap between specific and non-specific antibodies (**Figure S19C-F**), indicating some limitations for the Harvey method in predicting non-specificity of antibodies as compared to Nbs.

### 2.9 Prediction of non-specificity in antibody drug development programs

#### Consequences of non-specificity in the clinic

Non-specific binding of therapeutic antibodies can lead to significant adverse effects in the clinic. Such antibodies can bind to structurally unrelated off-targets and thereby potentially result in unwanted toxicity^57^ or reduced efficacy^23^. They can also interact with tissues like subcutis and thereby result in faster clearance via pinocytosis independently of FcRn^54,55,58^. Ultimately, non-specificity can compromise the safety and efficacy of therapeutic antibodies, potentially resulting in clinical trial failures and increased development costs. Thus, early-stage prediction of non-specificity, such as during selection and optimisation stage, is essential to reduce the risk of failing at late-stage during clinical trials. Otherwise, the further into the development program, the harder it becomes to allow additional protein engineering to mitigate biophysical liabilities, as new *in vitro* and *in vivo* data must be reproduced.

A powerful strategy to address this problem is to combine *in silico* prediction with *in vitro* developability assessment. To identify and flag non-specific antibodies, we propose a combined strategy that integrates *in silico* prediction models with traditional *in vitro* developability assessments during the lead optimization stage in the drug development process. This hybrid approach leverages the strengths of computational predictions and experimental validations, ensuring the selection of lead candidates with high developability potential.

## 3. Conclusions

In this study, we developed ML models to predict the non-specificity of antibodies, utilizing both PLMs and biophysical properties to embed antibody sequences. In agreement with previous reports on different datasets^59^, our results indicate that the VH domain, particularly the H-CDR loops, as the main contributor to non-specificity, and that the biophysical parameter pI is a key biophysical driver of non-specificity. The resulting computational models enable the prediction of non-specific interactions of antibodies with accuracy of 71% in 10-fold CV, thus providing a valuable tool to guide the design and selection of monoclonal antibodies with improved specificity and efficacy. These findings have important implications for the development of safe and efficacious lead candidates with high developability potential.

## 4. Methods

### 4.1 Data sources

All antibody datasets used in this study were retrieved from public sources, as well as upon request from Prof. Debora Marks and Prof. Andrew Kruse (Harvey dataset), and Dr. Tushar Jain and Dr. Dane Wittrup (Jain dataset). A list of the datasets and their corresponding sources are reported Table 2.

**Table 2.** List of public antibody datasets with their corresponding size, non-specificity assay and reference.

| Dataset | Size | Poly-reactive assay | Reference |
| --- | --- | --- | --- |
| Boughter dataset | >1000 antibodies (HIV-1 broadly neutralizing, Influenza reactive, IgA mouse) of varying degree of non-specificity | ELISA with a panel of 7 ligands (DNA, insulin, LPS, albumin, cardiolipin, flagellin, KLH) | <sup>38</sup> |
| Jain dataset | 137 clinical stage IgG1-formatted antibodies | ELISA with a panel of 6 ligands (ssDNA, dsDNA, insulin, LPS, cardiolipin, KLH) | <sup>15</sup> |
| Shehata dataset | 398 antibodies from naïve, IgG memory, IgM memory and long-lived plasma cells | Poly-specific reagent (PSR) assay | <sup>51</sup> |
| Harvey dataset | >140 000 naïve nanobodies | Poly-specific reagent (PSR) assay | <sup>39</sup> |

### 4.2 Python programming

All coding was performed in Python using Spyder IDE and Jupyter Notebook (Anaconda software distribution)^60^, and a list of used Python modules is reported in **Table 3**.

**Table 3.** List of software and Python modules.

| Module | Additional information | Reference |
| --- | --- | --- |
| NumPy | https://numpy.org | <sup>61</sup> |
| SciPy | https://www.scipy.org | <sup>62</sup> |
| Statsmodels | https://www.statsmodels.org | <sup>63</sup> |
| Pandas | https://pandas.pydata.org | <sup>64</sup> |
| Matplotlib | https://matplotlib.org | <sup>65</sup> |
| Seaborn | https://seaborn.pydata.org | <sup>66</sup> |
| Scikit-Learn | https://scikit-learn.org | <sup>67</sup> |
| Json | https://docs.python.org/3/library/json | <sup>68</sup> |
| Collections | https://docs.python.org/3/library/collections | <sup>69</sup> |
| Itertools | https://docs.python.org/3/library/itertools | <sup>70</sup> |
| ANARCI | https://github.com/oxpig/ANARCI | <sup>71</sup> |

**Table 4.**
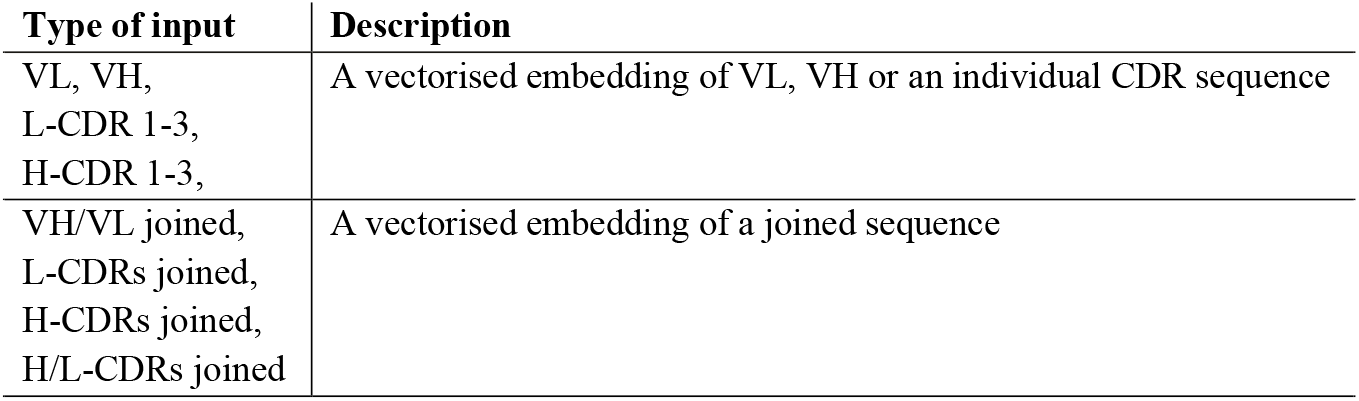
Type and description of sequence input for the binary classification models.

### 4.3 Training and validation of binary classification models

First, the Boughter dataset was parsed into three groups as previously done in ^38^: specific group (0 flags), mildly poly-reactive group (1-3 flags) and poly-reactive group (>3 flags). The primary sequences were annotated in the CDRs using ANARCI following the IMGT numbering scheme. Following this, 16 different antibody fragment sequences were assembled and embedded by three state-of-the-art protein language models (PLMs), ESM 1v^52^, Protbert bfd^53^, and AbLang2^44^, for representation of the physico-chemical properties and secondary/tertiary structure (**Table 4**). For the embeddings from the PLMs, *mean* (average of all token vectors) was used. The vectorised embeddings were served as features for training of binary classification models (e.g. LogisticReg, RandomForest, GaussianProcess, GradeintBoosting and SVM algorithms) for non-specificity (class 0: specific group, and class 1: poly-reactive group). The mildly poly-reactive group was left out from the training of the models.

The trained classification models were validated by (i) 3, 5 and 10-Fold cross-validation (CV), (ii) Leave-One Family-Out validation, e.g. training on HIV and Influenza reactive antibodies, while testing on mouse IgA antibodies, (iii) comparing probability of predicted poly-reactive class to true class, and (iv) testing on the Jain dataset. The evaluation metrics included accuracy, sensitivity and specificity (Eqs. 1-3).

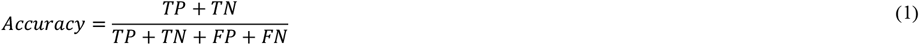

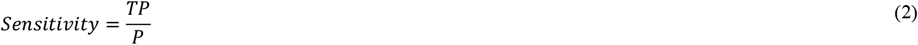

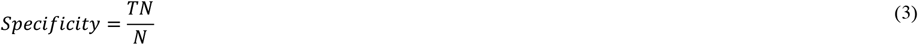

Where TP is true positive (true poly-reactive), TN is true negative (true specific), FP is false positive (false poly-reactive), FN is false negative (false specific), P is all positives (all poly-reactive), and N is all negatives (all specific).

### 4.4 Data availability

Data and models used in this publication are available at our GitHub repository: https://github.com/NovoNordisk-OpenSource/ML-predictions-of-Antibody-Non-Specificity.

## Acknowledgements

We would like to thank Prof. Debora Marks and Prof. Andrew Kruse for kindly sharing the dataset on 140 000 Nb clones assessed by the PSR assay from ^39^, as well as Dr. Tushar Jain and Dr. Dane Wittrup for kindly sharing the dataset on individual ELISA readouts from ^15^. We would also like to thank Mauricio Augilar Rangel, Dillon Rinauro and Ross Taylor from the University of Cambridge for their valuable discussions to this work. ChatGPT-5 mini (OpenAI) is acknowledged for use as a tool to draft the figure legends of this paper.

## Funding

Financial support from Novo Nordisk A/S is acknowledged.

P.S. is a Royal Society University Research Fellow (grant no. URF\R1\201461) and acknowledges funding from UK Research and Innovation (UKRI), and Engineering and Physical Sciences Research Council (grant no. EP/X024733/1).

The authors declare no competing financial interest.

## Notes

### Competing Interest Statement

The authors have declared no competing interest.

### Summary of Updates

The manuscript have been revised, references updated and link to access data and models have been included.

