## Supplementary material for "Prediction of Antibody Non-Specificity using Protein Language Models and Biophysical Parameters": SI

**Supporting Information**

Laila I. Sakhnini[1,3,*], Ludovica Beltrame[2], Simone Fulle[2,], Pietro Sormanni[3], Anette Henriksen[1], Nikolai Lorenzen[1], Michele Vendruscolo[3,*], Daniele Granata[2,*]

[1]Therapeutics Discovery, Novo Nordisk A/S, Copenhagen, Denmark
[2]Digital Chemistry and Design, Novo Nordisk A/S, Copenhagen, Denmark
[3]Centre for Misfolding Diseases, Department of Chemistry, University of Cambridge, UK

[*]Corresponding authors:

L.I. Sakhnini, M. Vendruscolo, D. Granata

### 18 List of Figures


| Figure | Description |
|---|---|
| S1 | Distribution of non-specificity from three public antibody datasets: (A) Boughter dataset, (B) UMAP projection of the sequence similarity distances of the H/L-CDRs for the antibodies within the Boughter dataset, (C) boxplot of H-CDR3 length of the antibodies within the Boughter dataset, (D) Jain dataset, (E) Shehata dataset, and (F) balance of non-specificity class within individual datasets. |
| S2 | 10-fold CV for different antibody domain input embedded by ESM 1v: panel (A) Accuracy, panel (B) Sensitivity, and panel (C) Specificity. |
| S3 | 10-fold CV for different antibody domain input embedded by ESM 1b: panel (A) Accuracy, panel (B) Sensitivity, and panel (C) Specificity. |
| S4 | 10-fold CV for different antibody domain input embedded by ESM 2: panel (A) Accuracy, panel (B) Sensitivity, and panel (C) Specificity. |
| S5 | 10-fold CV for different antibody domain input embedded by Protbert bfd: panel (A) Accuracy, panel (B) Sensitivity, and panel (C) Specificity. |
| S6 | 10-fold CV for different antibody domain input embedded by AntiBERTy: panel (A) Accuracy, panel (B) Sensitivity, and panel (C) Specificity. |
| S7 | 10-fold CV for different antibody domain input embedded by AbLang2: panel (A) Accuracy, panel (B) Sensitivity, and panel (C) Specificity. |
| S8 | 10-fold CV for different antibody domain input embedded by 68 different sequence-based descriptors: panel (A) Accuracy, panel (B) Sensitivity, and panel (C) Specificity. |
| S9 | Spearman correlation matrix of 68 VH-based sequence-based descriptors. |
| S10 | Bar plots showcasing accuracy of models with top descriptor combinations and their frequency among the top 15 models: panel (A) top 2-descriptor models (top 15 out of 300 combinations), panel (B) top 3-descriptor models (top 15 out of 2300 combinations), panel (C) top 4-descriptor models (top 15 out of 12650 combinations), and panel (D) top 5-descriptor models (top 15 out of 53130 combinations). Each bar represents a different combination of 2-5 descriptors, with error bars indicating the standard deviation of 10-fold CV accuracy. The red line and red shaded area indicate the 10-fold CV accuracy and standard deviation, respectively, for ESM 1v VH-based LogisticReg model. |
| S11 | Visualisation of feature importance (absolute value of Eigenvalues) for 5 components from the PCA of VH sequence descriptors. |
| S12 | VH-based LogisticReg models showcasing validation performance (k-Fold CV and Leave-One-Family-Out) for PLM- and descriptor-based models: (A) Sensitivity and (B) Specificity. Bar plot for accuracy can be found in Figure 2C in main paper. |
| S13 | Boxplot showing the predicted non-specificity probabilities for the respective ELISA flag of the Jain dataset using VH-based Logistic Regression (ESM 1v). The boxplot displays the median, interquartile range, and outliers, with strength of regression-like trend indicated by SCC and p-value (<0.001). |
| S14 | Confusion matrices for VH-based LogisticReg models across different datasets showcasing the number of antibodies correctly and incorrectly predicted as specific (label 0) and non-specific (label 1). Each panel represents a different model and dataset combination: panel (A) ESM 1v VH-based LogisticReg model tested on the Jain dataset, panel (B) Top 5-descriptors VH-based LogisticReg model tested on the Jain dataset, panel (C) ESM 1v VH-based LogisticReg model tested on the Shehata dataset, panel (D) Top 5-descriptors VH-based LogisticReg model tested on the Shehata dataset, panel (E) ESM 1v VH-based LogisticReg model tested on the Harvey dataset, and panel (F) Top 5-descriptors VH-based LogisticReg model tested on the Harvey dataset. |

| | |
|---|---|
| S15 | Distribution of top VH-based sequence descriptors for specific and non-specific antibodies in the Boughter dataset (mildly non-specific antibodies excluded). Histograms and boxplots show the distribution of various descriptors for specific (blue) and non-specific (red) antibodies. Each subplot represents a different descriptor: panel (A) theoretical pI, panel (B) bulkiness, panel (C) disorder propensity according to DisProt, panel (D) percentage of accessible residues, panel (E) Aggrescan av4, panel (F) average flexibility using BP scale, panel (G) beta turn propensity according to Chou-Fasman, panel (H) high-performance liquid chromatography retention using HFBA, and panel (I) number of hotspots according to Aggrescan. For each descriptor, the top boxplots display the distribution for specific and non-specific antibodies, with the median and interquartile range indicated. The histograms illustrate the frequency of antibodies within different value bins, providing insights into the characteristics of each descriptor. |
| S16 | Distribution of top VH-based sequence descriptors for specific and non-specific antibodies in the Jain dataset (mildly non-specific antibodies excluded). Histograms and boxplots show the distribution of various descriptors for specific (blue) and non-specific (red) antibodies. Each subplot represents a different descriptor: panel (A) theoretical pI, panel (B) bulkiness, panel (C) disorder propensity according to DisProt, panel (D) percentage of accessible residues, panel (E) Aggrescan av4, panel (F) average flexibility using BP scale, panel (G) beta turn propensity according to Chou-Fasman, panel (H) high-performance liquid chromatography retention using HFBA, and panel (I) number of hotspots according to Aggrescan. For each descriptor, the top boxplots display the distribution for specific and non-specific antibodies, with the median and interquartile range indicated. The histograms illustrate the frequency of antibodies within different value bins, providing insights into the characteristics of each descriptor. |
| S17 | Distribution of top VH-based sequence descriptors for specific and non-specific antibodies in the Shehata dataset. Histograms and boxplots show the distribution of various descriptors for specific (blue) and non-specific (red) antibodies. Each subplot represents a different descriptor: panel (A) theoretical pI, panel (B) bulkiness, panel (C) disorder propensity according to DisProt, panel (D) percentage of accessible residues, panel (E) Aggrescan av4, panel (F) average flexibility using BP scale, panel (G) beta turn propensity according to Chou-Fasman, panel (H) high-performance liquid chromatography retention using HFBA, and panel (I) number of hotspots according to Aggrescan. For each descriptor, the top boxplots display the distribution for specific and non-specific antibodies, with the median and interquartile range indicated. The histograms illustrate the frequency of antibodies within different value bins, providing insights into the characteristics of each descriptor. |
| S18 | Distribution of top VH-based sequence descriptors for specific and non-specific antibodies in the Harvey dataset. Histograms and boxplots show the distribution of various descriptors for specific (blue) and non-specific (red) antibodies. Each subplot represents a different descriptor: panel (A) theoretical pI, panel (B) bulkiness, panel (C) disorder propensity according to DisProt, panel (D) percentage of accessible residues, panel (E) Aggrescan av4, panel (F) average flexibility using BP scale, panel (G) beta turn propensity according to Chou-Fasman, panel (H) high-performance liquid chromatography retention using HFBA, and panel (I) number of hotspots according to Aggrescan. For each descriptor, the top boxplots display the distribution for specific and non-specific antibodies, with the median and interquartile range indicated. The histograms illustrate the frequency of antibodies within different value bins, providing insights into the characteristics of each descriptor. |
| S19 | Performance of Harvey *et al.* predictor (2022) on various datasets: panels (A-B) Boughter dataset, panels (C-D) Jain dataset, and panels (E-F) Shehata dataset. |

### List of Tables


| Table | Description |
|---|---|
| S1 | Overview of biophysical descriptors. All the descriptors are derived from Schrödinger but those marked (*), which have been calculated with the Biopython ProteinAnalysis module. Further documentation on the Schrödinger descriptors can be found at https://support.schrodinger.com/s/article/827119. |
| S2 | Summary of descriptor importance and model performance for a VH-based LogisticReg model trained on all descriptors (excluding charge at pH 6 and 7.4). For each descriptor, the following is provided; (i) cluster number assigned by hierarchical clustering of Spearman correlation coefficients as a way to group redundant descriptors, (ii) LogisticReg coefficients from the logistic regression model, (ii) absolute value of LogisticReg coefficients from the LogisticReg model, (iii) the decrease in model accuracy when the descriptor is permuted, indicating its importance (based on 10-fold CV on test data), (iv) the model accuracy when the specific descriptor is left out, indicating its unique contribution, and (v) the accuracy of the model using only the single descriptor. Descriptors are listed along with their respective cluster number and their importance metrics, highlighting their contribution to the model's performance and their individual predictive power. |

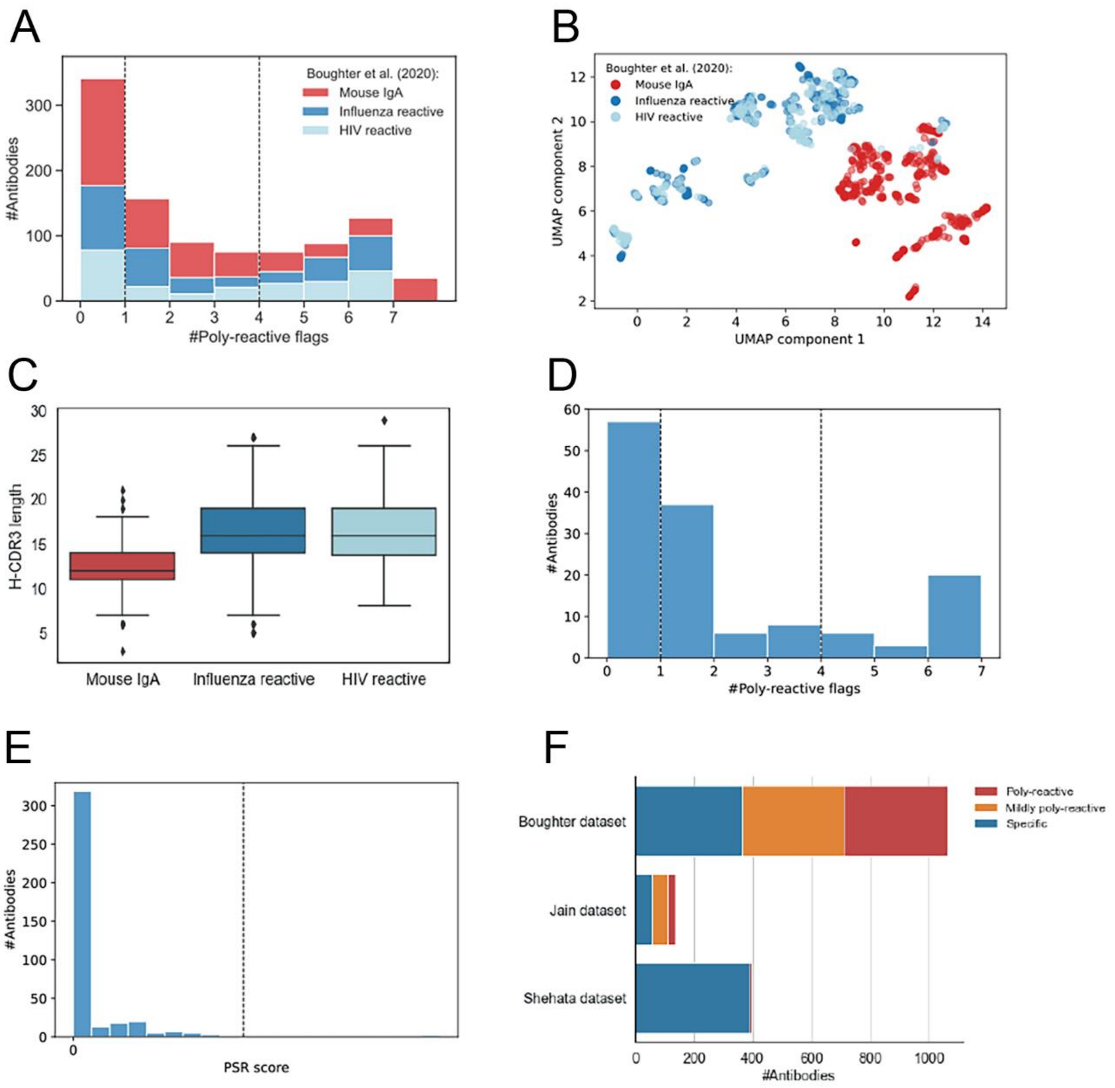


**Figure S1.** Distribution of non-specificity from three public antibody datasets: (A) Boughter dataset, (B) UMAP projection of the sequence similarity distances of the H/L-CDRs for the antibodies within the Boughter dataset, (C) boxplot of H-CDR3 length of the antibodies within the Boughter dataset, (D) Jain dataset, (E) Shehata dataset, and (F) balance of non-specificity class within individual datasets.

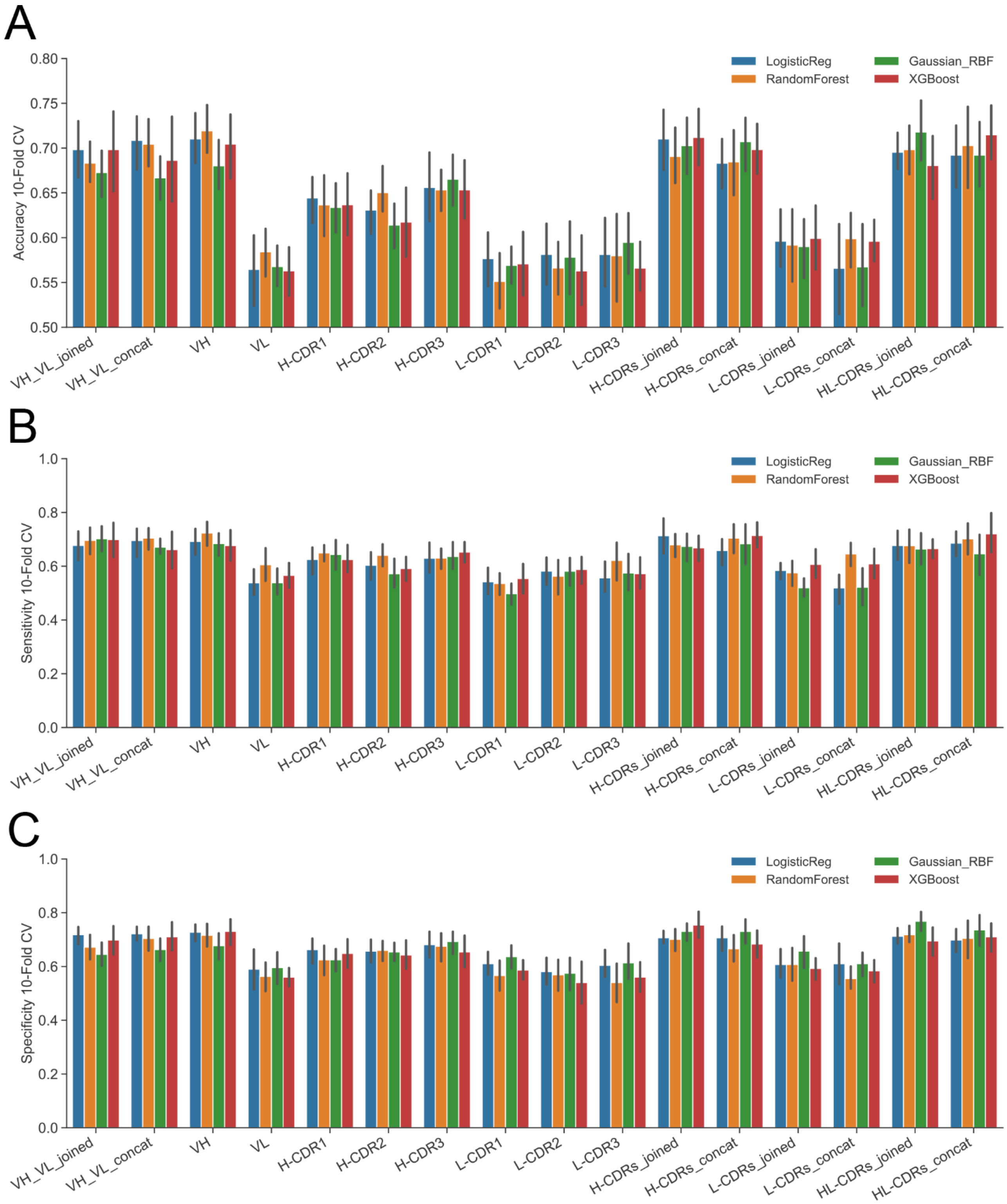


**Figure S2.** 10-fold CV for different antibody domain input embedded by ESM 1v: (A) Accuracy, (B) Sensitivity, and (C) Specificity.

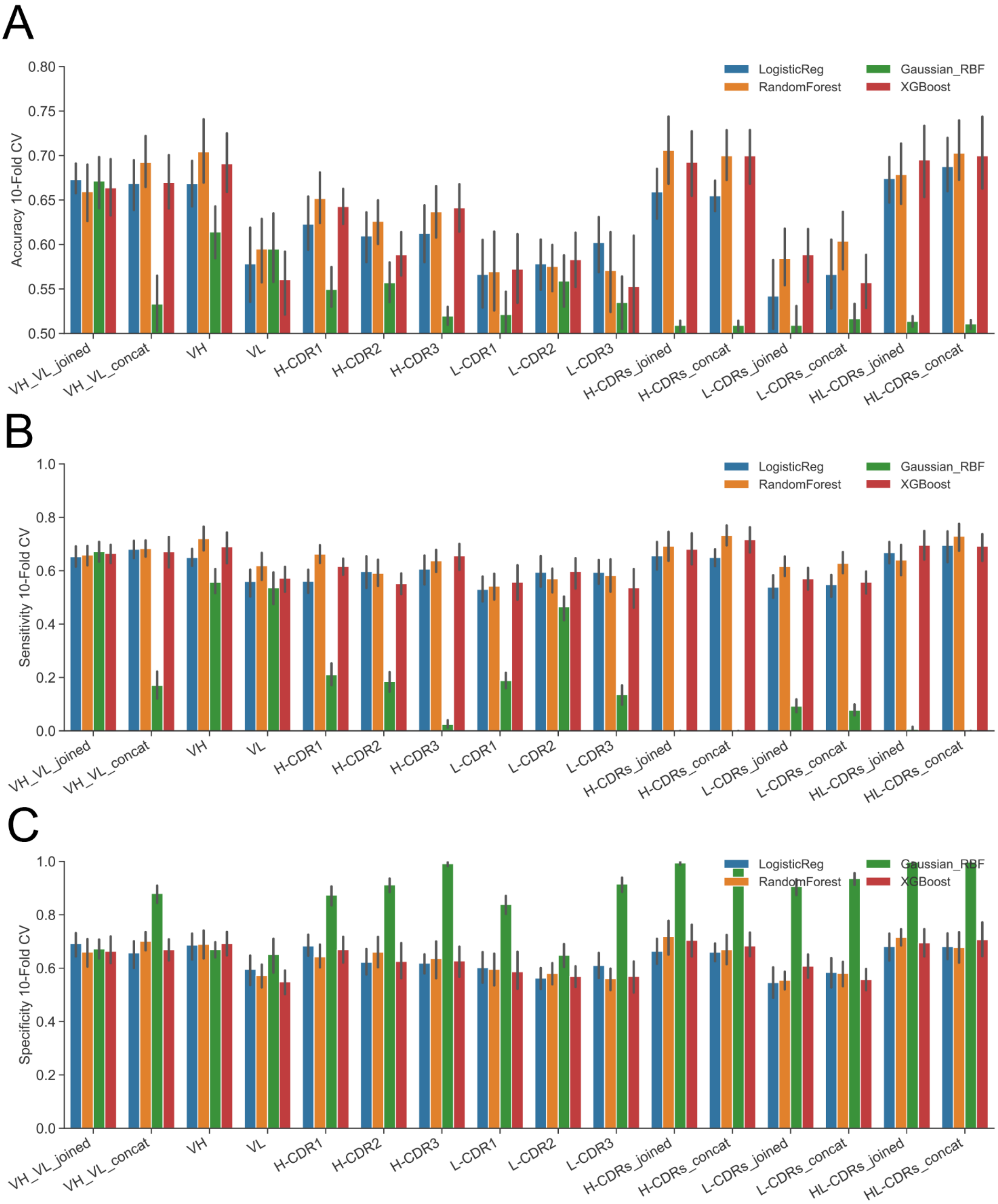


**Figure S3.** 10-fold CV for different antibody domain input embedded by ESM 1b: (A) Accuracy, (B) Sensitivity, and (C) Specificity.

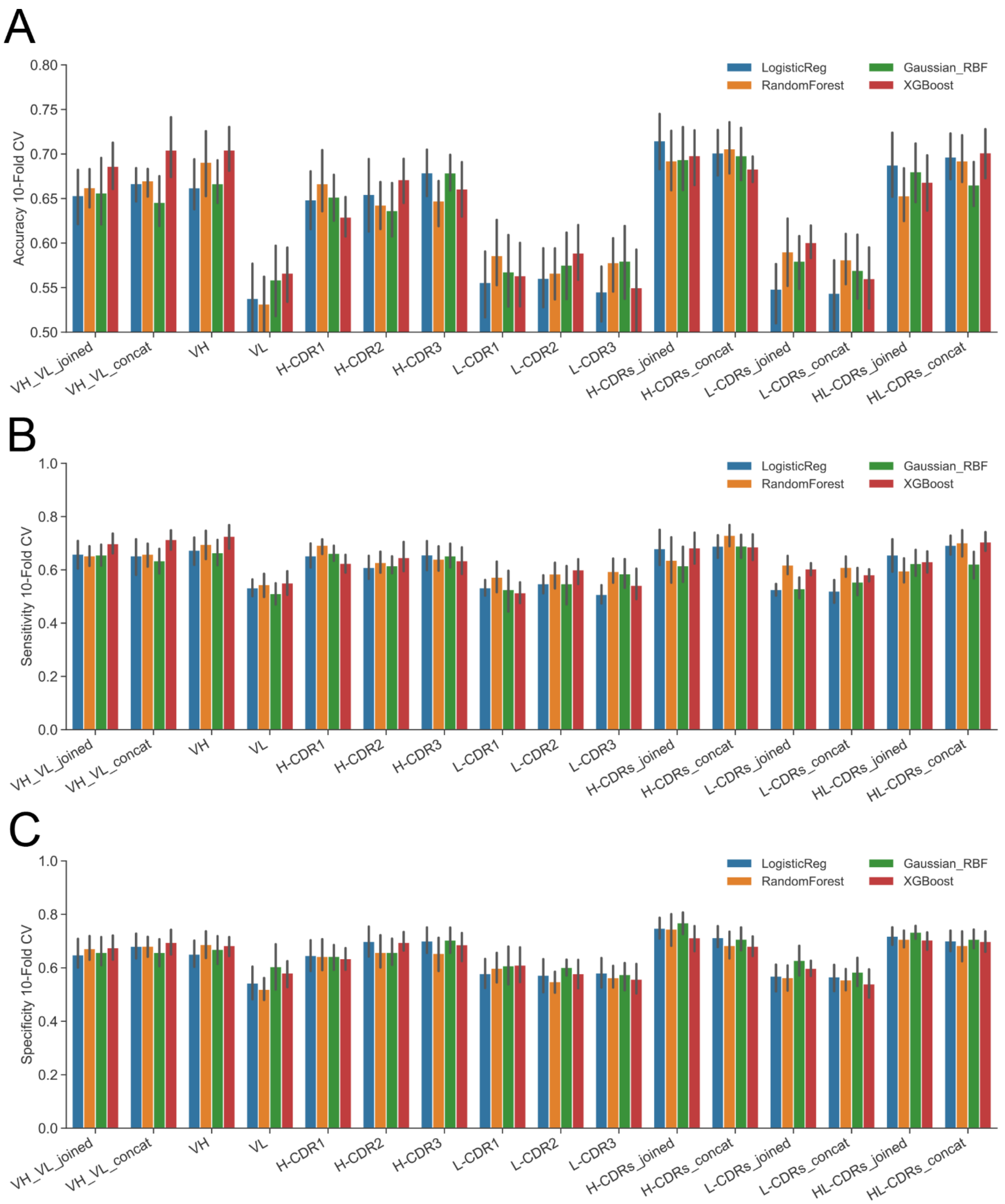


**Figure S4.** 10-fold CV for different antibody domain input embedded by ESM 2: (A) Accuracy, (B) Sensitivity, and (C) Specificity.

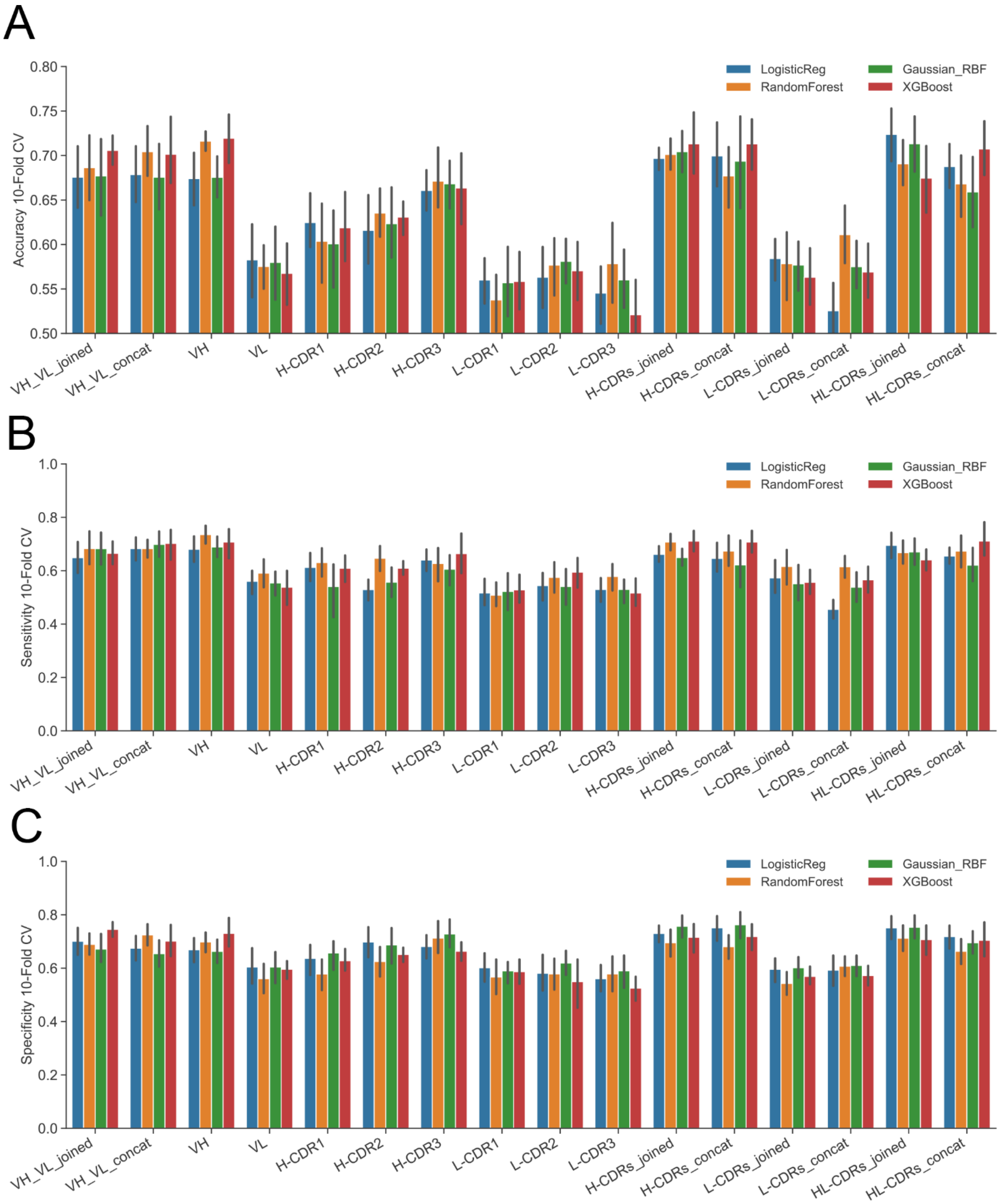


**Figure S5.** 10-fold CV for different antibody domain input embedded by Protbert bfd: (A) Accuracy, (B) Sensitivity, and (C) Specificity.

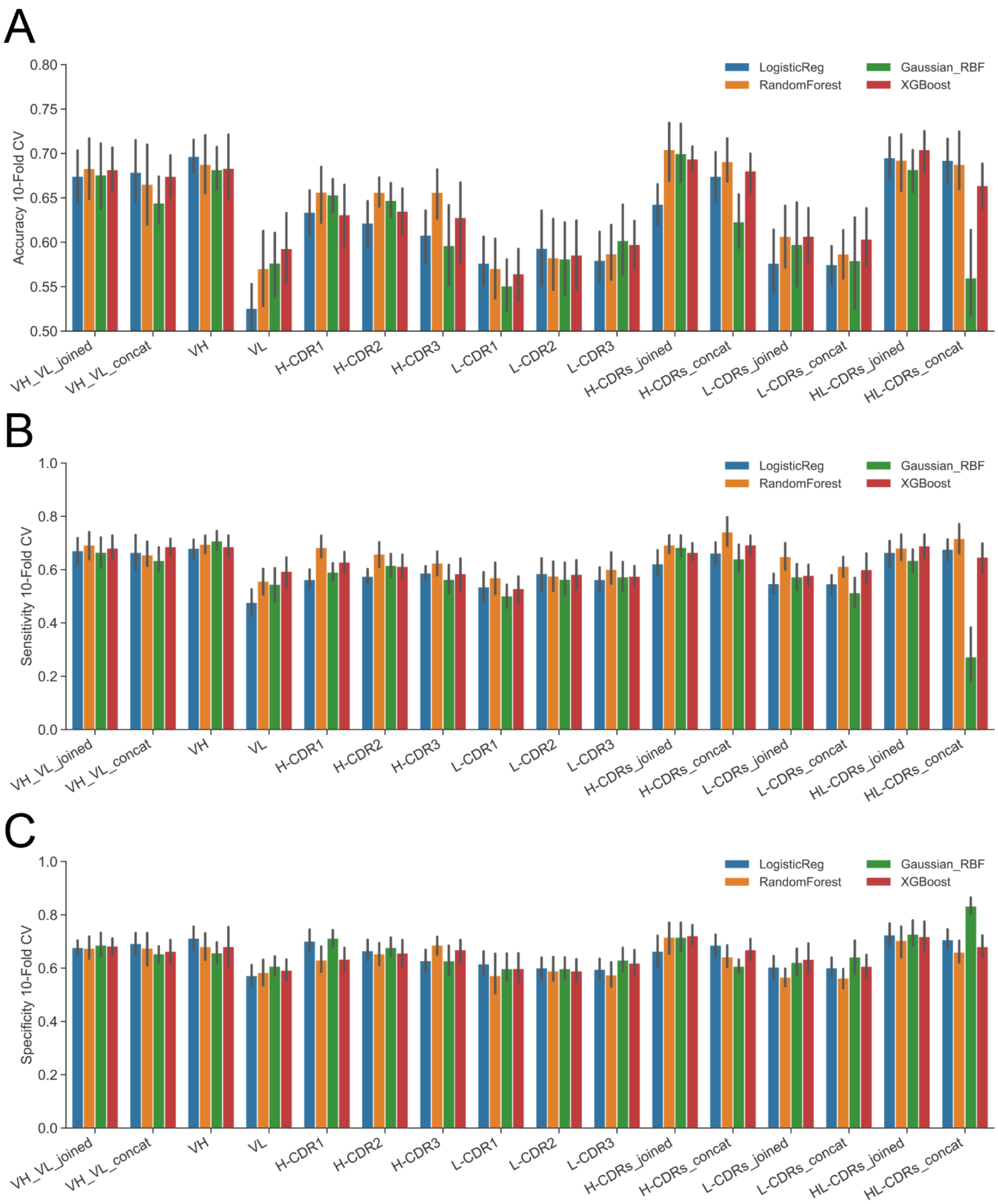


**Figure S6.** 10-fold CV for different antibody domain input embedded by AntiBERTy: (A) Accuracy, (B) Sensitivity, and (C) Specificity.

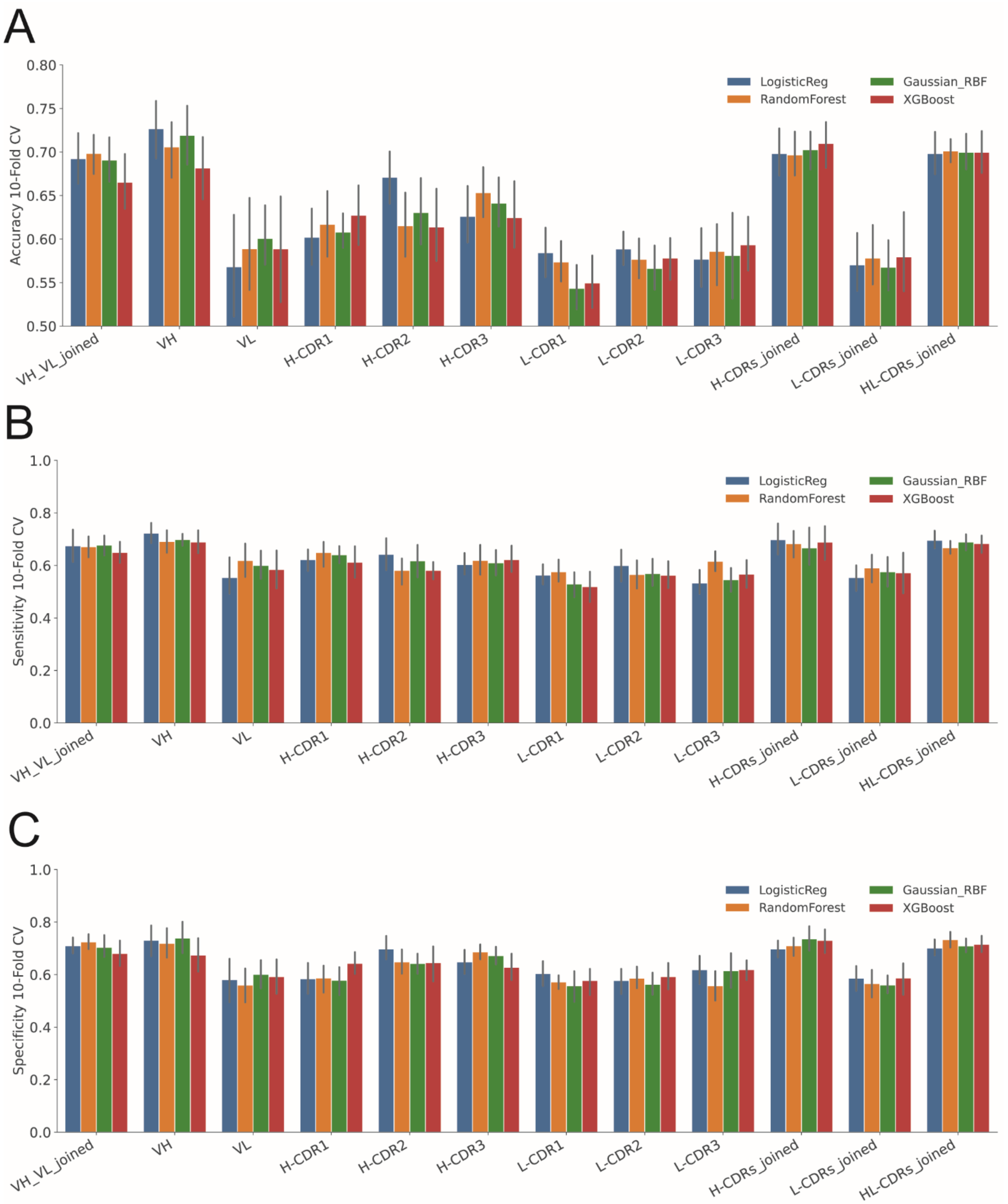


**Figure S7.** 10-fold CV for different antibody domain input embedded by AbLang2: (A) Accuracy, (B) Sensitivity, and (C) Specificity.

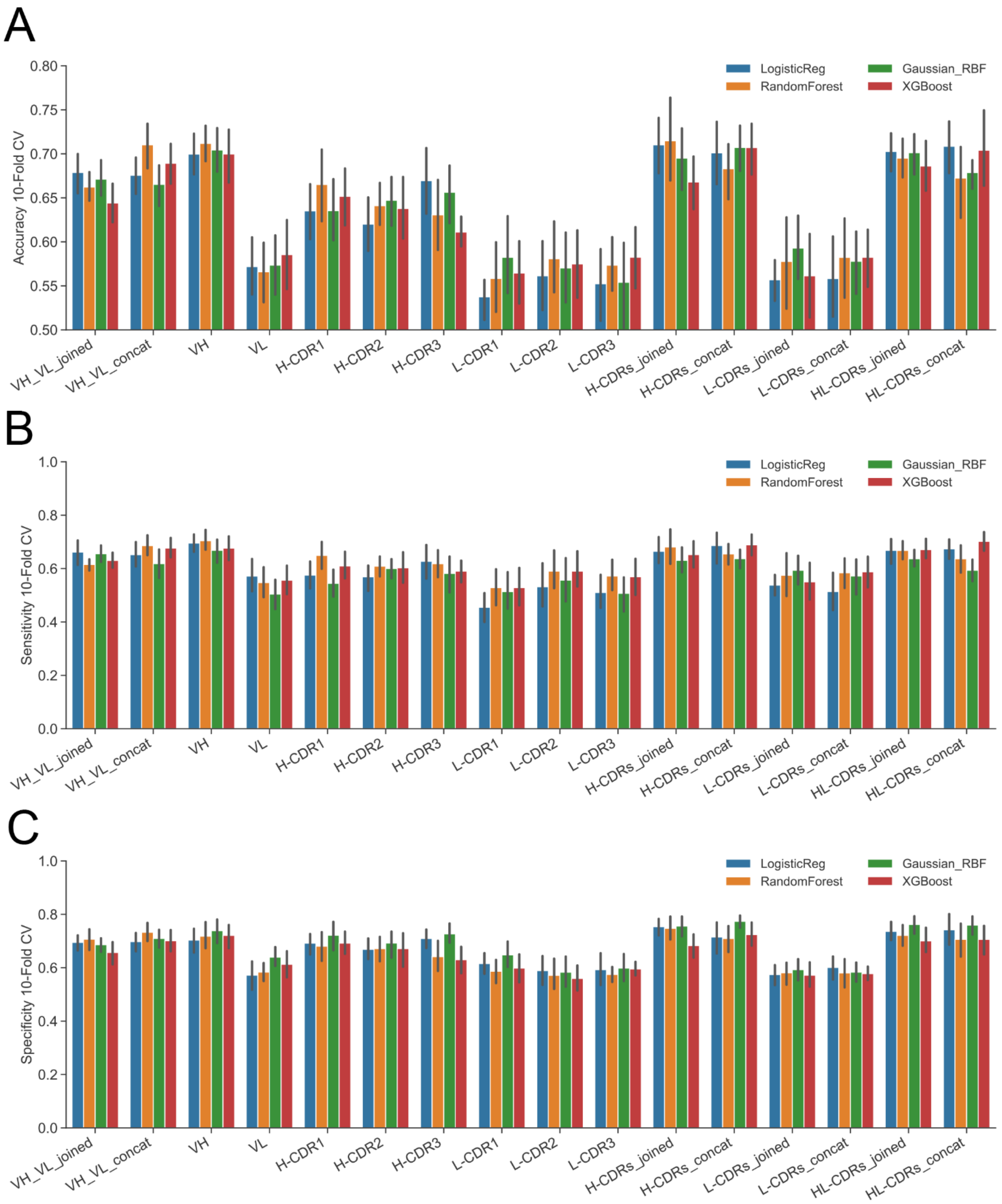


**Figure S8.** 10-fold CV for different antibody domain input embedded by 68 different sequence-based descriptors: (A) Accuracy, (B) Sensitivity, and (C) Specificity.

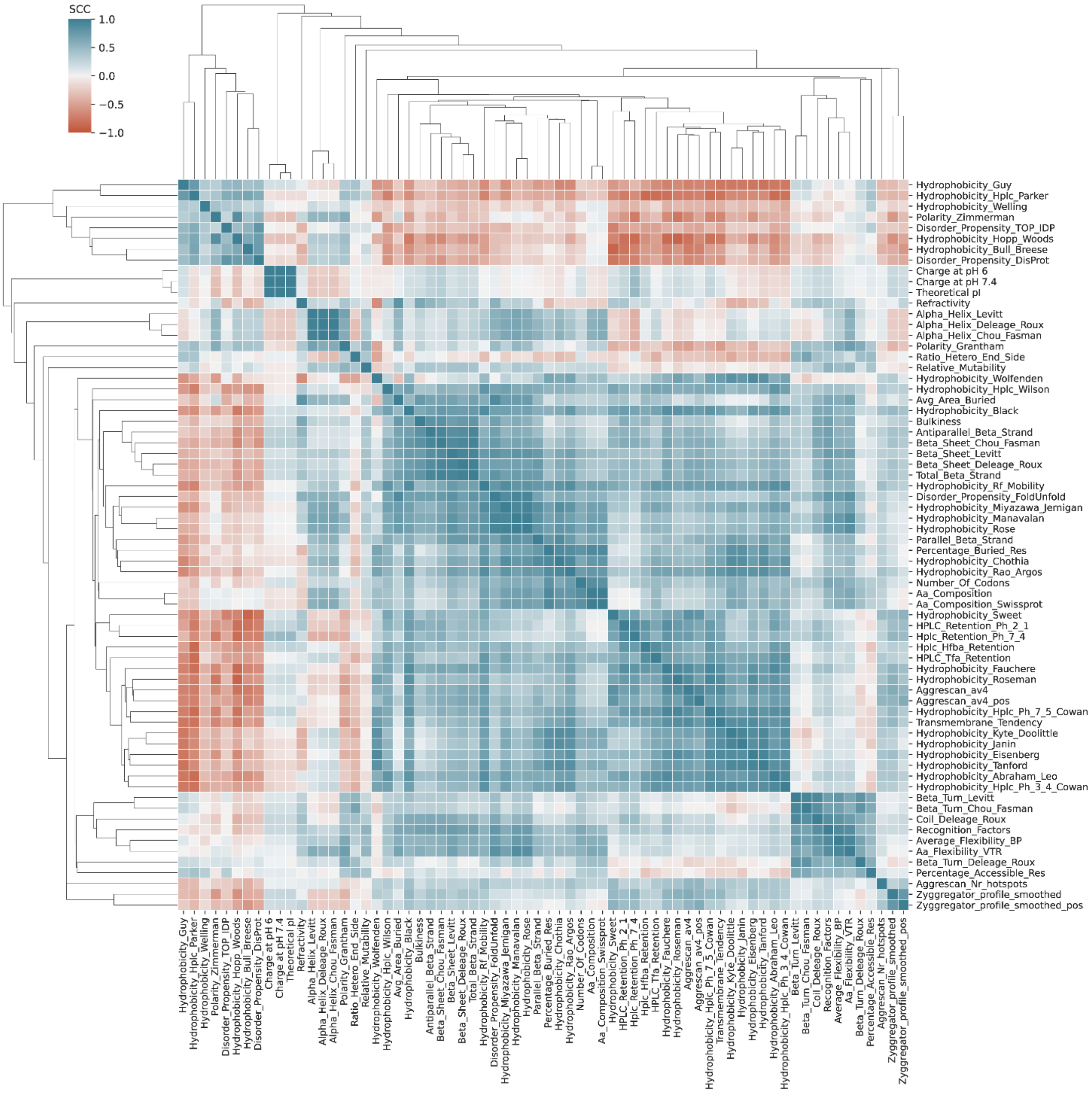


**Figure S9.** Spearman correlation matrix of 68 VH-based sequence-based descriptors.

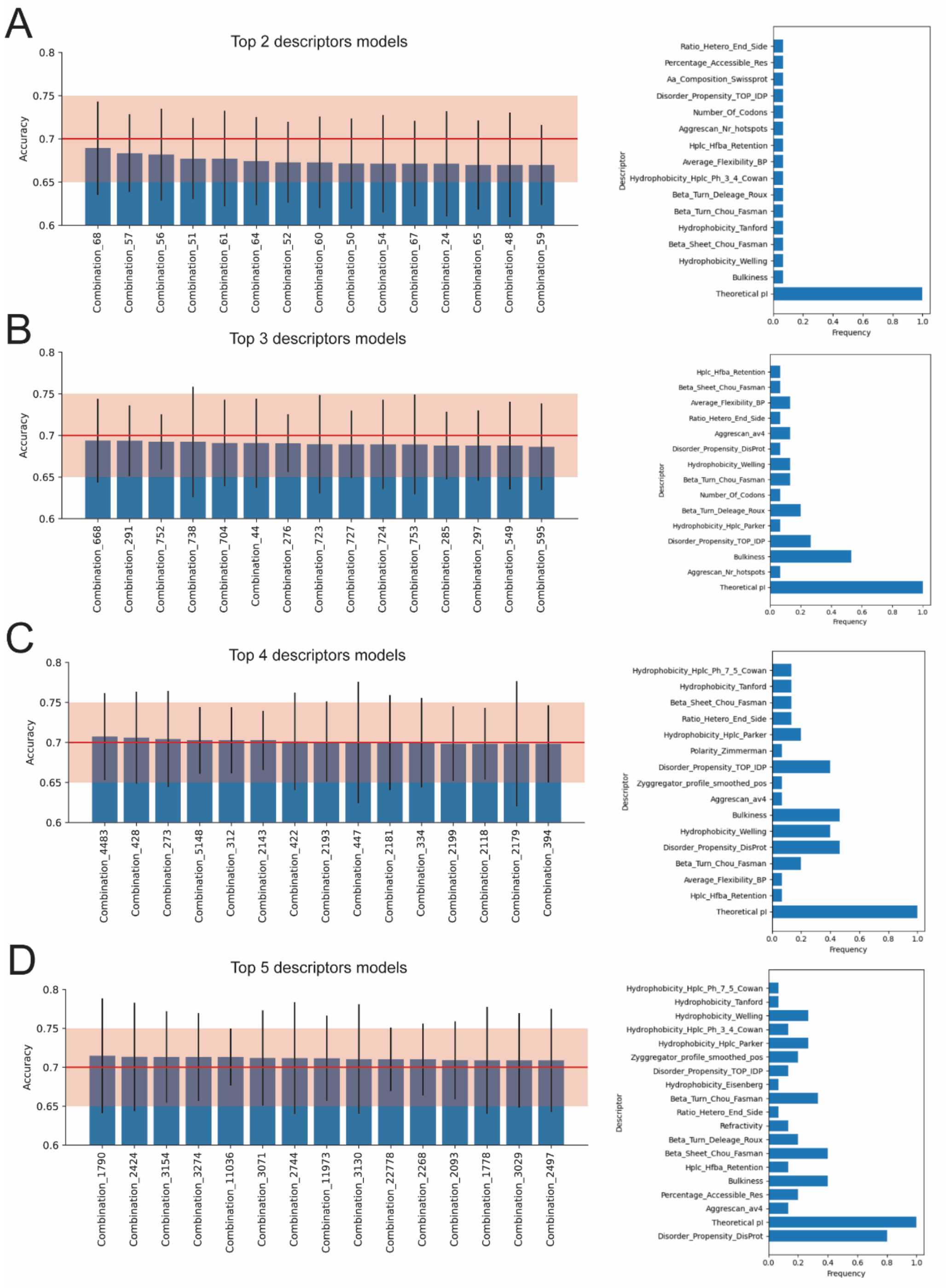


**Figure S10.** Bar plots showcasing accuracy of models with top descriptor combinations and their frequency among the top 15 models: (A) top 2-descriptor models (top 15 out of 300 combinations), (B) top 3-descriptor models (top 15 out of 2300 combinations), (C) top 4-descriptor models (top 15 out of 12650 combinations), and (D) top 5-descriptor models (top 15 out of 53130 combinations). Each bar represents a different combination of 2-5 descriptors, with error bars indicating the standard deviation of 10-fold CV accuracy. The red line and red shaded area indicate the 10-fold CV accuracy and standard deviation, respectively, for ESM 1v VH-based LogisticReg model.

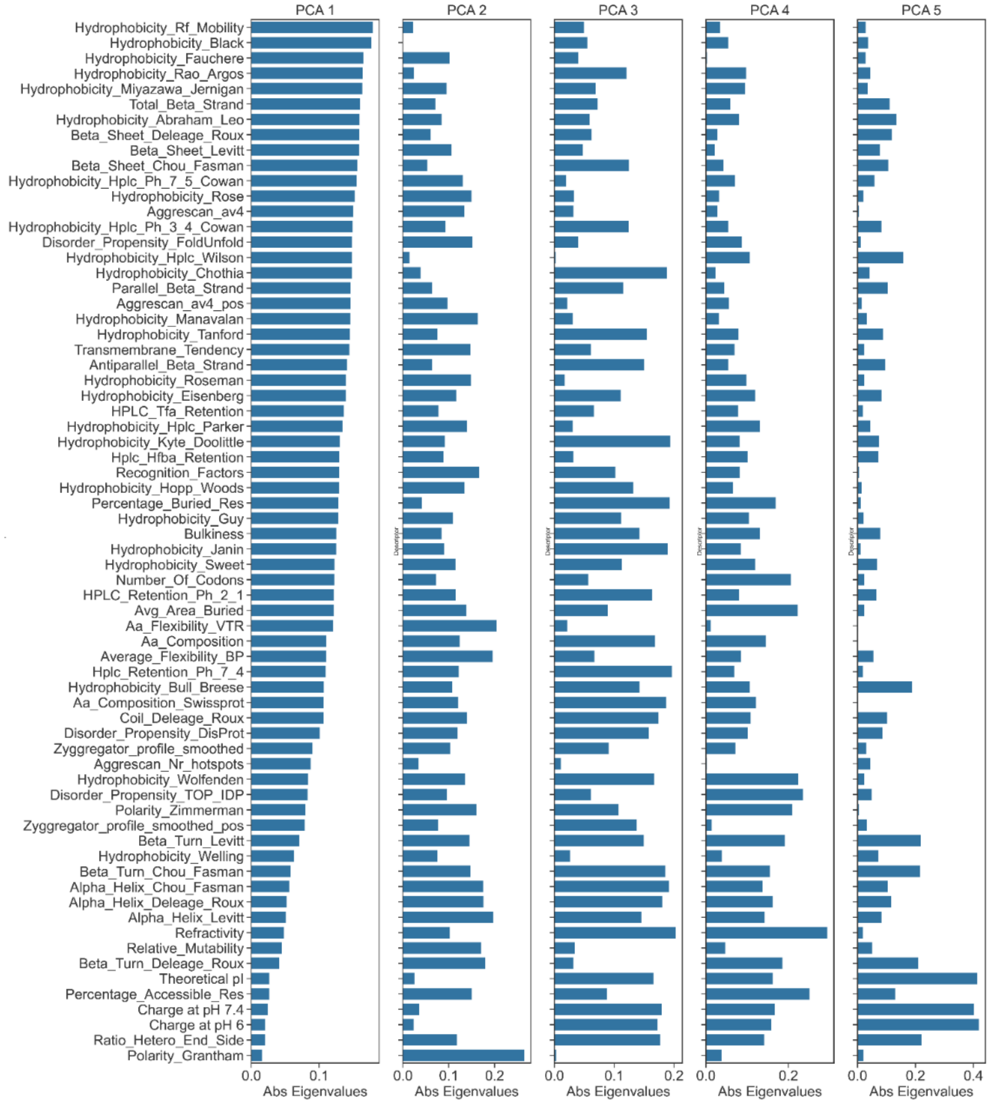


**Figure S11.** Visualisation of feature importance (absolute value of Eigenvalues) for 5 components from the PCA of VH sequence descriptors.

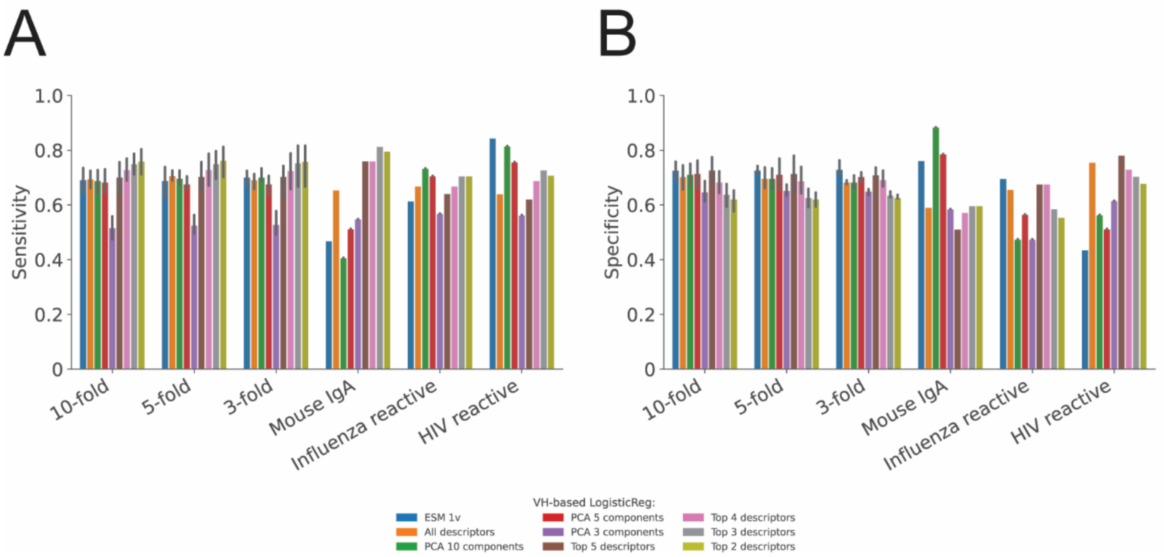


**Figure S12.** VH-based LogisticReg models showcasing validation performance (k-Fold CV and Leave-One-Family-Out) for PLM- and descriptor-based models: (A) Sensitivity and (B) Specificity. Bar plot for accuracy can be found in Figure 2C in main paper.

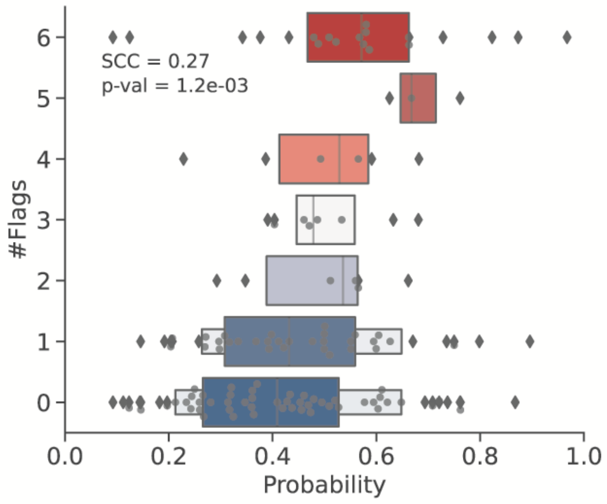


**Figure S13.** Boxplot showing the predicted non-specificity probabilities for the respective ELISA flag of the Jain dataset using VH-based Logistic Regression (ESM 1v). The boxplot displays the median, interquartile range, and outliers, with strength of regression-like trend indicated by SCC and p-value (<0.001).

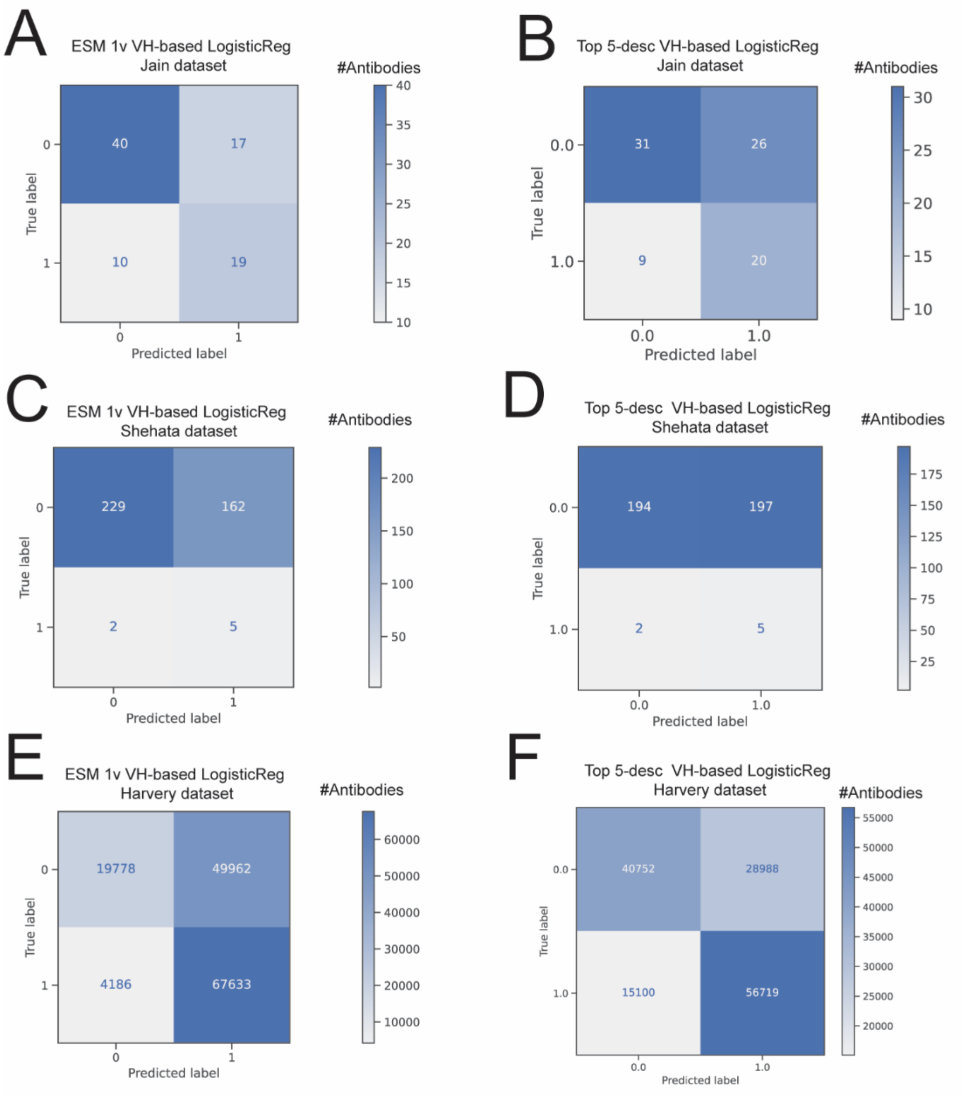


**Figure S14.** Confusion matrices for VH-based LogisticReg models across different datasets showcasing the number of antibodies correctly and incorrectly predicted as specific (label 0) and non-specific (label 1). Each panel represents a different model and dataset combination: panel (A) ESM 1v VH-based LogisticReg model tested on the Jain dataset, panel (B) Top 5-descriptors VH-based LogisticReg model tested on the Jain dataset, panel (C) ESM 1v VH-based LogisticReg model tested on the Shehata dataset, panel (D) Top 5-descriptors VH-based LogisticReg model tested on the Shehata dataset, panel (E) ESM 1v VH-based LogisticReg model tested on the Harvey dataset, and panel (F) Top 5-descriptors VH-based LogisticReg model tested on the Harvey dataset.

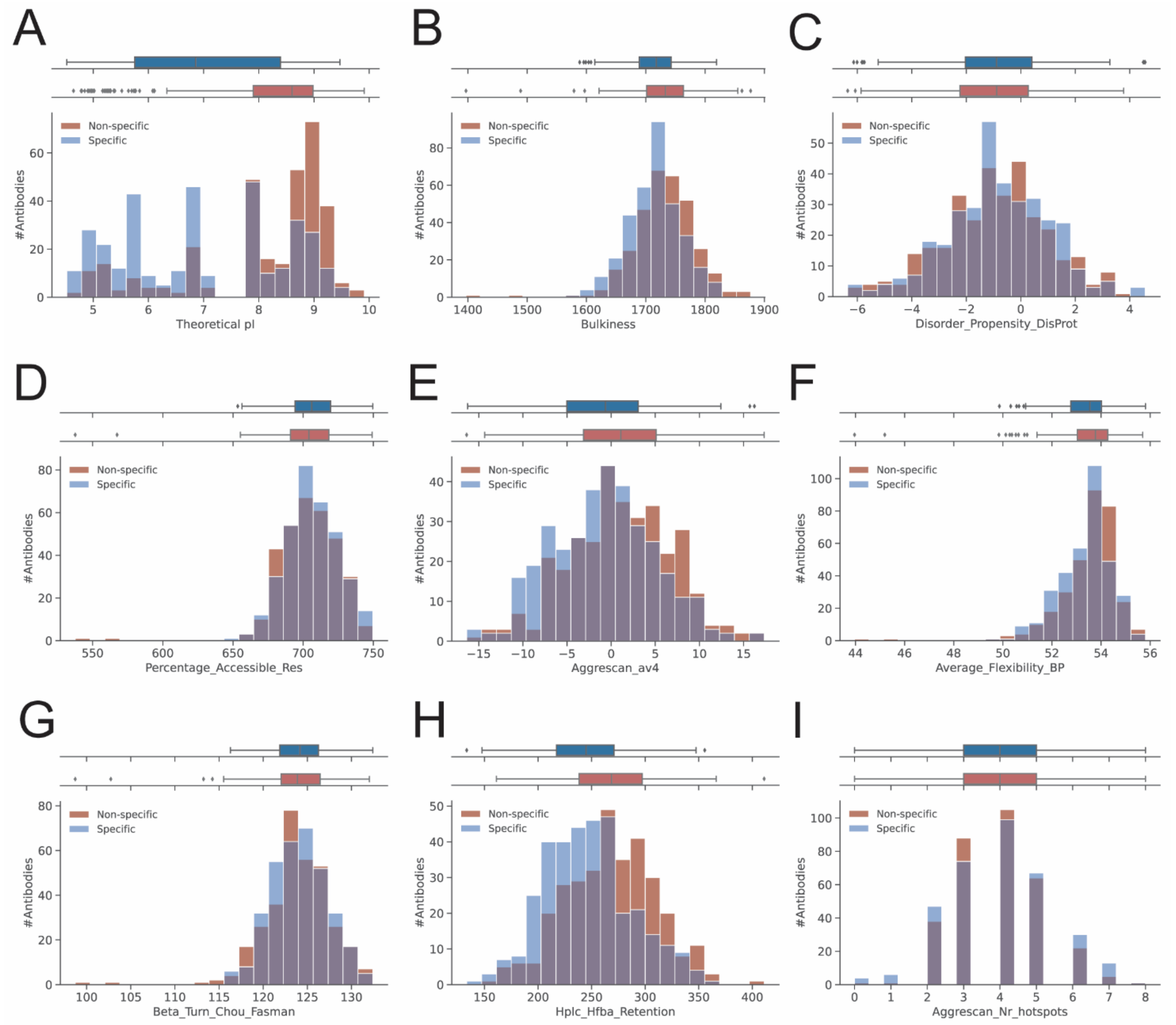


**Figure S15. Boughter dataset.** Distribution of top VH-based sequence descriptors for specific and non-specific antibodies (mildly non-specific antibodies excluded). Histograms and boxplots show the distribution of various descriptors for specific (blue) and non-specific (red) antibodies. Each subplot represents a different descriptor: panel (A) theoretical pI, panel (B) bulkiness, panel (C) disorder propensity according to DisProt, panel (D) percentage of accessible residues, panel (E) Aggrescan av4, panel (F) average flexibility using BP scale, panel (G) beta turn propensity according to Chou-Fasman, panel (H) high-performance liquid chromatography retention using HFBA, and panel (I) number of hotspots according to Aggrescan. For each descriptor, the top boxplots display the distribution for specific and non-specific antibodies, with the median and interquartile range indicated. The histograms illustrate the frequency of antibodies within different value bins, providing insights into the characteristics of each descriptor.

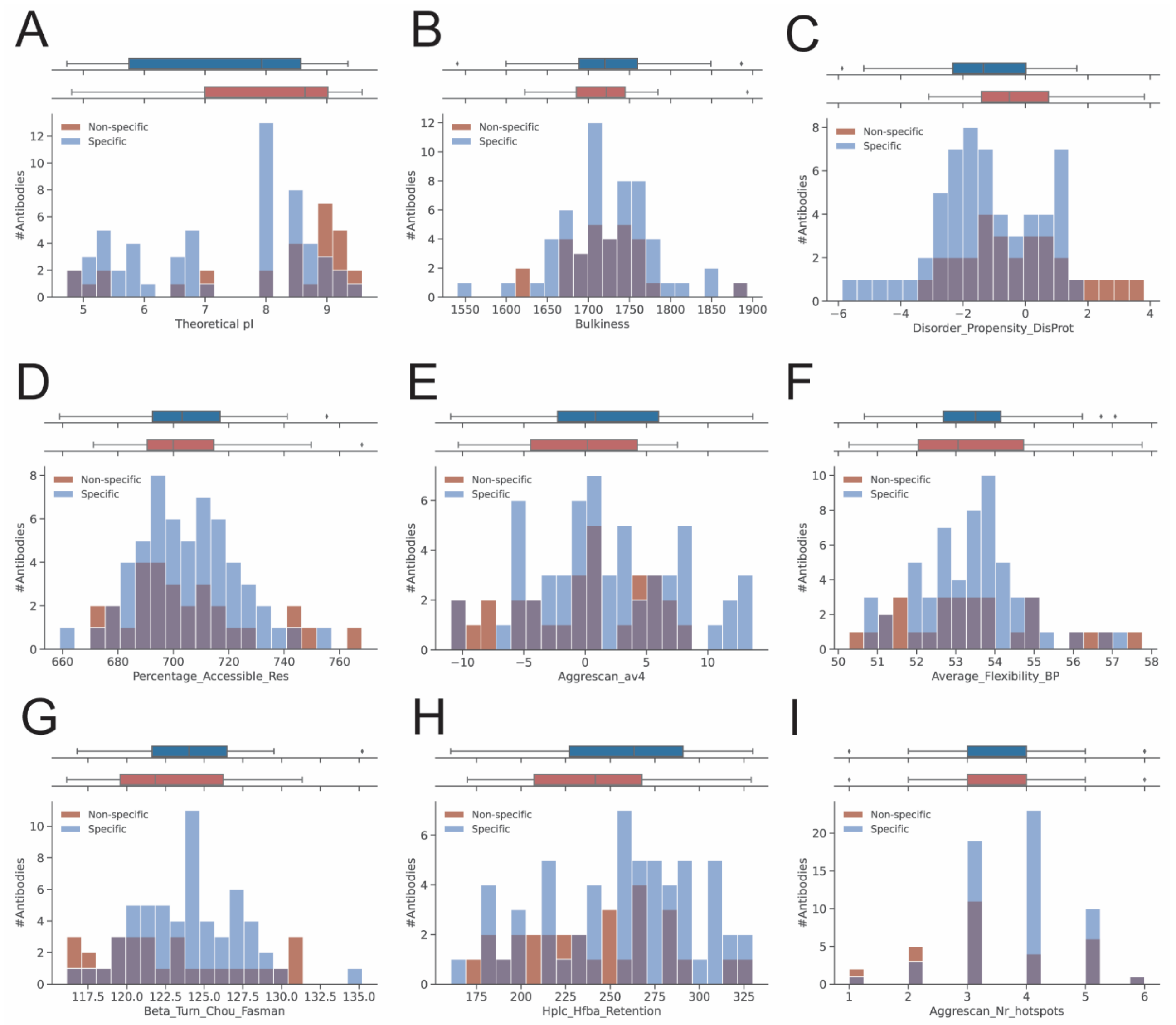


**Figure S16. Jain dataset.** Distribution of top VH-based sequence descriptors for specific and non-specific antibodies (mildly non-specific antibodies excluded). Histograms and boxplots show the distribution of various descriptors for specific (blue) and non-specific (red) antibodies. Each subplot represents a different descriptor: panel (A) theoretical pI, panel (B) bulkiness, panel (C) disorder propensity according to DisProt, panel (D) percentage of accessible residues, panel (E) Aggrescan av4, panel (F) average flexibility using BP scale, panel (G) beta turn propensity according to Chou-Fasman, panel (H) high-performance liquid chromatography retention using HFBA, and panel (I) number of hotspots according to Aggrescan. For each descriptor, the top boxplots display the distribution for specific and non-specific antibodies, with the median and interquartile range indicated. The histograms illustrate the frequency of antibodies within different value bins, providing insights into the characteristics of each descriptor.

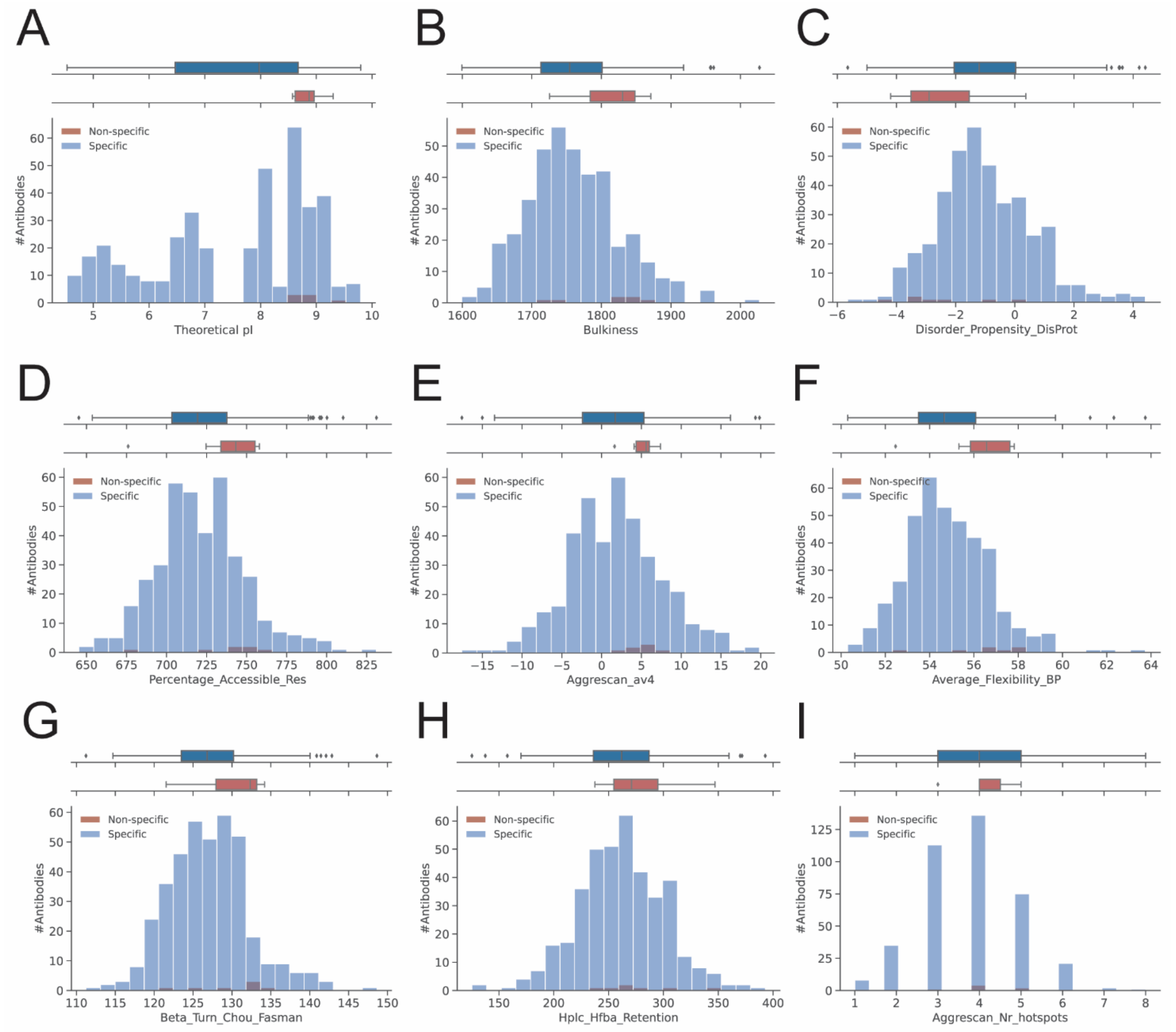


**Figure S17. Shehata dataset.** Distribution of top VH-based sequence descriptors for specific and non-specific antibodies. Histograms and boxplots show the distribution of various descriptors for specific (blue) and non-specific (red) antibodies. Each subplot represents a different descriptor: panel (A) theoretical pI, panel (B) bulkiness, panel (C) disorder propensity according to DisProt, panel (D) percentage of accessible residues, panel (E) Aggrescan av4, panel (F) average flexibility using BP scale, panel (G) beta turn propensity according to Chou-Fasman, panel (H) high-performance liquid chromatography retention using HFBA, and panel (I) number of hotspots according to Aggrescan. For each descriptor, the top boxplots display the distribution for specific and non-specific antibodies, with the median and interquartile range indicated. The histograms illustrate the frequency of antibodies within different value bins, providing insights into the characteristics of each descriptor.

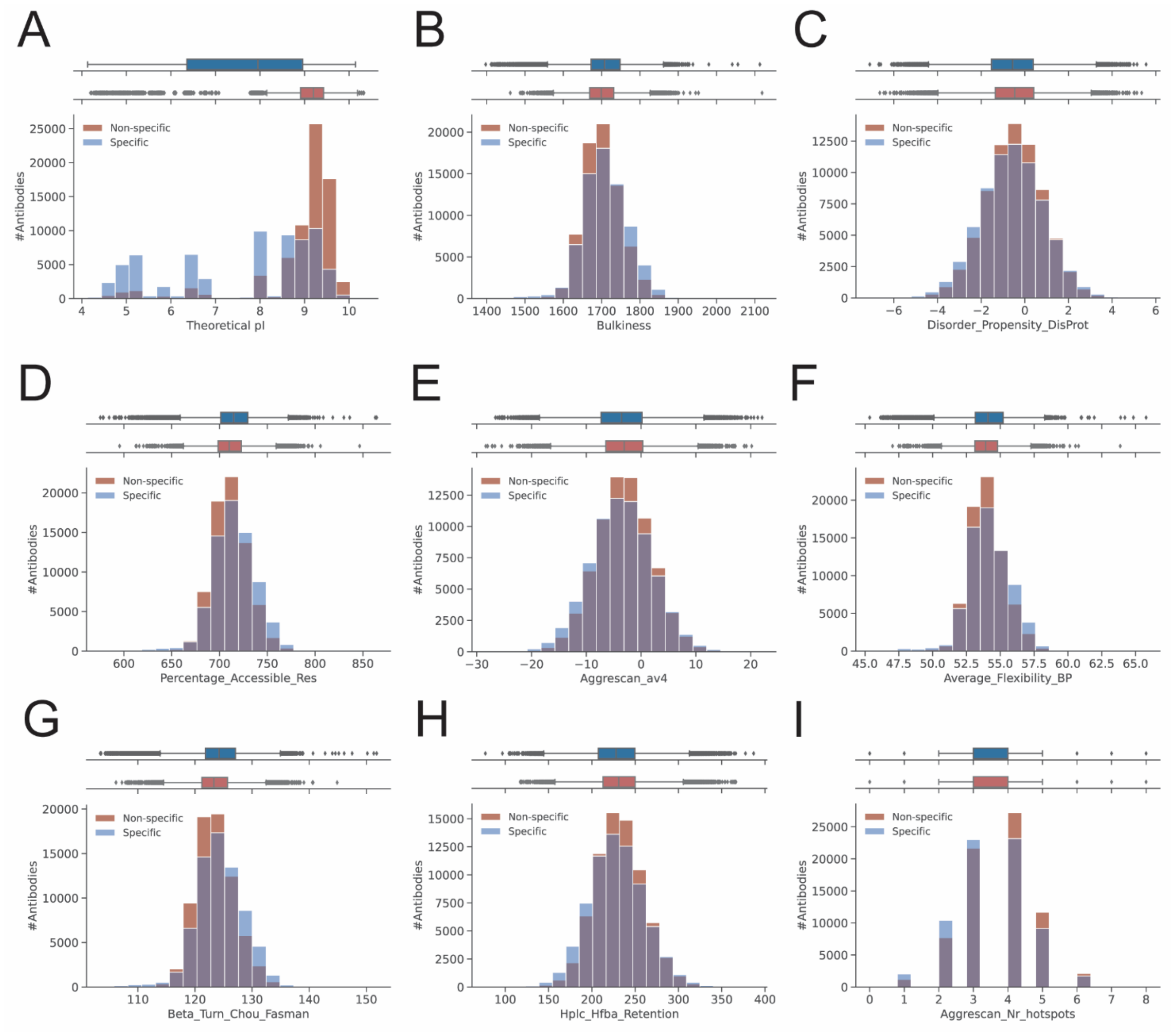


**Figure S18. Harvey dataset.** Distribution of top VH-based sequence descriptors for specific and non-specific antibodies. Histograms and boxplots show the distribution of various descriptors for specific (blue) and non-specific (red) antibodies. Each subplot represents a different descriptor: panel (A) theoretical pI, panel (B) bulkiness, panel (C) disorder propensity according to DisProt, panel (D) percentage of accessible residues, panel (E) Aggrescan av4, panel (F) average flexibility using BP scale, panel (G) beta turn propensity according to Chou-Fasman, panel (H) high-performance liquid chromatography retention using HFBA, and panel (I) number of hotspots according to Aggrescan. For each descriptor, the top boxplots display the distribution for specific and non-specific antibodies, with the median and interquartile range indicated. The histograms illustrate the frequency of antibodies within different value bins, providing insights into the characteristics of each descriptor.

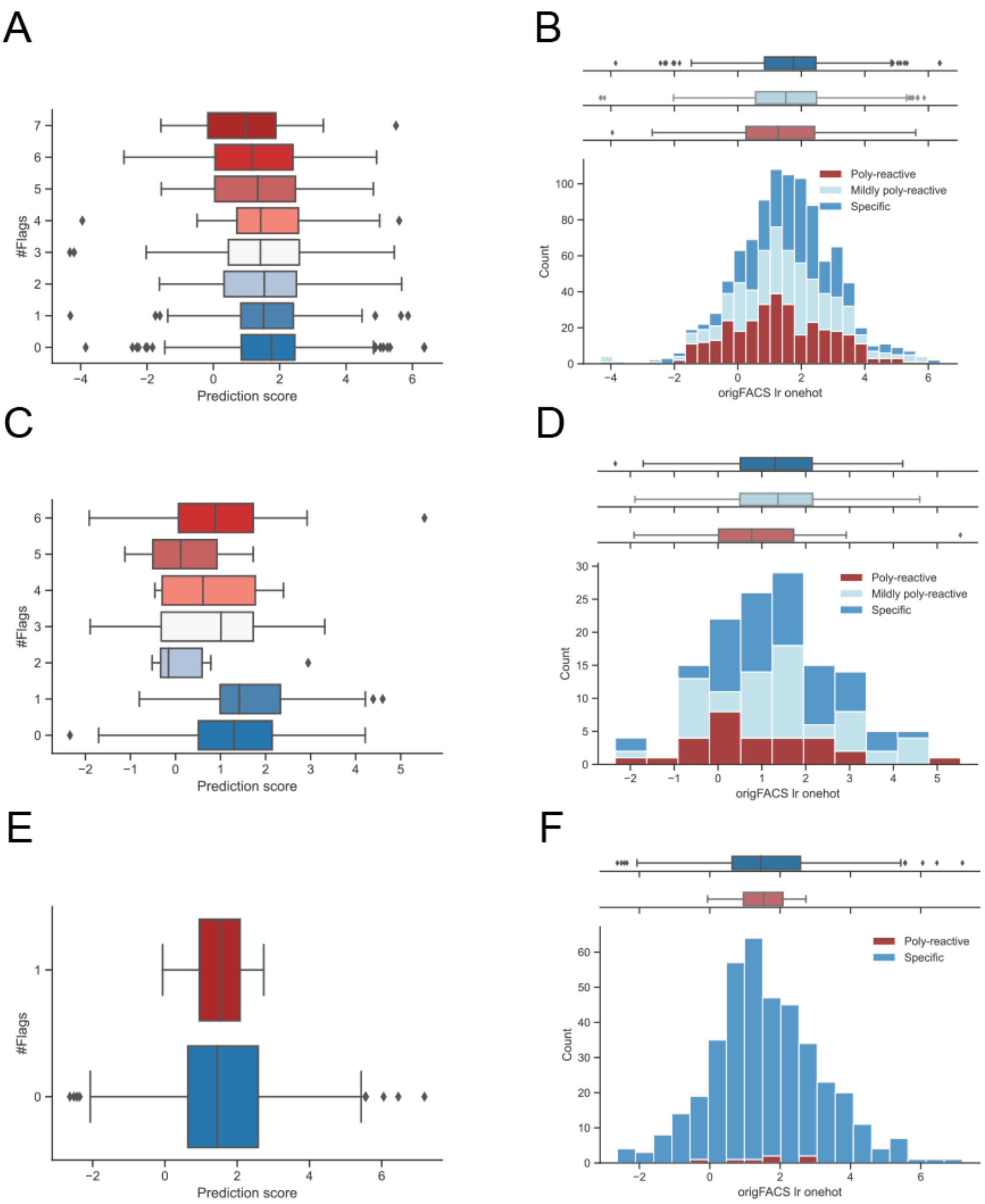


**Figure S19.** Performance of Harvey *et al.* predictor (2022): (A-B) Boughter dataset, (C-D) Jain dataset, and (E-F) Shehata dataset.

**Table S1**. Overview of biophysical descriptors. All the descriptors are derived from Schrödinger but those marked (*), which have been calculated with the Biopython ProteinAnalysis module. Further documentation on the Schrödinger descriptors can be found at: https://support.schrodinger.com/s/article/827119.

| # | Descriptor | Definition |
|---|---|---|
| 1 | AGGRESCAN_Nr_hotspots | Number of aggregation hotspots computed by the Aggrescan algorithm (http://bioinf.uab.es/aap/aap_help.html) |
| 2 | Aa_Composition | The total of amino acid composition as described by McCaldon and Argos (Proteins: Structure, Function and Genetics 4:99-122(1988)) |
| 3 | Aa_Composition_Swissprot | The total value of amino acid composition based on SwissProt annotation (Release notes for UniProtKB/Swiss-Prot release 2013_04 - April 2013) |
| 4 | Aa_Flexibility_VTR | The total amino acid flexibility as defined by Vihinen, Torkkila, and Rikonen (https://www.ncbi.nlm.nih.gov/pubmed/8090708) |
| 5 | Aggrescan_av4 | a4v values over a sliding window, as determined by the Aggrescan algorithm |
| 6 | Aggrescan_av4_pos | a4v positive values over a sliding window, as determined by the Aggrescan algorithm |
| 7 | All_Aggrescan_a4v_pos | The sum of the average of a4v positive values over a sliding window, as determined by the Aggrescan algorithm |
| 8 | Alpha_Helix_Chou_Fasman | Alpha helix propensity, as defined by Chou and Fasman (Adv. Enzym. 47:45-148(1978)) |
| 9 | Alpha_Helix_Deleage_Roux | Alpha helix propensity, as defined by Deleage and Roux (Protein Engineering 1:289-294(1987)) |
| 10 | Alpha_Helix_Levitt | Alpha helix propensity, as defined by Levitt (Biochemistry 17:4277-4285(1978)) |
| 11 | Antiparallel_Beta_Strand | Antiparallel beta strand propensity, as defined by Lifson and Sander (Nature 282:109-111(1979)) |
| 12 | Average_Flexibility_BP | Total amino acid flexibility, as defined by Bhaskaran and Ponnusamy (Int. J. Pept. Protein. Res. 32:242-255(1988)) |
| 13 | Avg_Area_Buried | Average standard-state to folded-protein buried area, as defined by Rose et al. (Science 229:834-838(1985)) |
| 14 | Beta_Sheet_Chou_Fasman | Beta sheet propensity, as defined by Chou and Fasman (Adv. Enzym. 47:45-148(1978)) |
| 15 | Beta_Sheet_Deleage_Roux | Beta sheet propensity, as defined by Deleage and Roux (Protein Engineering 1:289-294(1987)) |
| 16 | Beta_Sheet_Levitt | Beta sheet propensity, as defined by Levitt (Biochemistry 17:4277-4285(1978)) |
| 17 | Beta_Turn_Chou_Fasman | Beta turn propensity, as defined by Chou and Fasman (Adv. Enzym. 47:45-148(1978)) |
| 18 | Beta_Turn_Deleage_Roux | Beta turn propensity, as defined by Deleage and Roux (Protein Engineering 1:289-294(1987)) |
| 19 | Beta_Turn_Levitt | Beta turn propensity, as defined by Levitt (Biochemistry 17:4277-4285(1978)) |
| 20 | Bulkiness | Total amino acid bulkiness (J. Theor. Biol. 21:170-201(1968)) |
| 21 | Charge at pH 6* | Charge of the protein at pH 6 |
| 22 | Charge at pH 7.4* | Charge of the protein at pH 7.4 |
| 23 | Coil_Deleage_Roux | Total score for coil, as defined by Deleage and Roux (Protein Engineering 1:289-294(1987)) |
| 24 | Disorder_Propensity_DisProt | Total disorder promotion propensity (https://www.ncbi.nlm.nih.gov/pubmed/17578581) |
| 25 | Disorder_Propensity_FoldUnfold | Total disorder promotion propensity (https://www.ncbi.nlm.nih.gov/pubmed/15498936) |
| 26 | Disorder_Propensity_TOP_IDP | Total disorder propensity for intrinsic disorder, based on the TOP-IDP scale model (https://www.ncbi.nlm.nih.gov/pmc/articles/PMC2676888/) |
| 27 | HPLC_Retention_Ph_2_1 | Total value of retention coefficients in HPLC at pH 2.1 (Proc. Natl. Acad. Sci. USA 77:1632-1636(1980)) |

| | | |
|---|---|---|
| 28 | HPLC_Tfa_Retention | Total value of retention coefficients in HPLC/TFA (Anal. Biochem. 124:201-208(1982)) |
| 29 | Hplc_Hfba_Retention | Total value of retention coefficients in HFBA (Anal. Biochem. 124:201-208(1982)) |
| 30 | Hplc_Retention_Ph_7_4 | Total value of retention coefficients in HPLC at pH 7.4 (Proc. Natl. Acad. Sci. USA 77:1632-1636(1980)) |
| 31 | Hydrophobicity_Abraham_Leo | Total hydrophobicity, as defined by Abraham and Leo (Proteins: Structure, Function and Genetics 2:130-152(1987)) |
| 32 | Hydrophobicity_Black | Total hydrophobicity, as definedy by Black (Anal. Biochem. 193:72-82(1991)) |
| 33 | Hydrophobicity_Bull_Breese | Total hydrophobicity, as definedy by Bull and Breese (Arch. Biochem. Biophys. 161:665-670(1974)) |
| 34 | Hydrophobicity_Chothia | Total hydrophobicity based on proportion of buried residues (95%), as defined by Chothia (J. Mol. Biol. 105:1-14(1976)) |
| 35 | Hydrophobicity_Eisenberg | Total normalized consensus hydrophobicity, as defined by Eisenberg et al. (J. Mol. Biol. 179:125-142(1984)) |
| 36 | Hydrophobicity_Fauchere | Total hydrophobicity, as defined by Fauchere (Eur. J. Med. Chem. 18:369-375(1983)) |
| 37 | Hydrophobicity_Guy | Total hydrophobicity based on free energy of transfer, as defined by Guy (Biophys J. 47:61-70(1985)) |
| 38 | Hydrophobicity_Hopp_Woods | Total hydrophilicity, as defined by Hopp & Woods (Proc. Natl. Acad. Sci. U.S.A. 78:3824-3828(1981)) |
| 39 | Hydrophobicity_Hplc_Parker | Total hydrophilicity derived from HPLC peptide retention times, as defined by Parker et al. (Biochemistry 25:5425-5431(1986)) |
| 40 | Hydrophobicity_Hplc_Ph_3_4_Cowan | Total hydrophobicity determined by HPLC at ph 3.4 , as defined by Cowan and Whittaker (Peptide Research 3:75-80(1990)) |
| 41 | Hydrophobicity_Hplc_Ph_7_5_Cowan | Total hydrophobicity determined by HPLC at ph 7.5, as defined by Cowan and Whittaker (Peptide Research 3:75-80(1990)) |
| 42 | Hydrophobicity_Hplc_Wilson | Total hydrophobicity derived from HPLC peptide retention times, as defined by Wilson et al. (Biochem. J. 199:31-41(1981)) |
| 43 | Hydrophobicity_Janin | Total hydrophobicity based on dG of transfer from inside to outside of a globular protein, as defined by Janin (Nature 277:491-492(1979)) |
| 44 | Hydrophobicity_Kyte_Doolittle | Total hydrophobicity, as defined by Kyte and Dolittle (J. Mol. Biol. 157:105-132(1982)) |
| 45 | Hydrophobicity_Manavalan | Total average surrounding hydrophobicity, as defined by Manavalan and Ponnusamy (Nature 275:673-674(1978)) |
| 46 | Hydrophobicity_Miyazawa_Jernigan | Hydrophobicity, as defined by Miyazawa and Jernigan (Macromolecules 18:534-552(1985)) |
| 47 | Hydrophobicity_Rao_Argos | Total transmembrane helix parameters, as defined by Rao and Argos (Biochim. Biophys. Acta 869:197-214(1986)) |
| 48 | Hydrophobicity_Rf_Mobility | Total hydrophobicity based on chromatograohic mobility, as defined by Aboderin (Int. J. Biochem. 2:537-544(1971)) |
| 49 | Hydrophobicity_Rose | Total hydrophobicity based on mean fractional exposed area loss (average area buried/standard state area), as defined by Rose (Science 229:834-838(1985)) |
| 50 | Hydrophobicity_Roseman | Total hydrophobicity, as defined by Roseman (J. Mol. Biol. 200:513-522(1988)) |
| 51 | Hydrophobicity_Sweet | Total optimized matching hydrophobicity, as defined by Sweet (J. Mol. Biol. 171:479-488(1983)) |
| 52 | Hydrophobicity_Tanford | Total hydrophobicity, as defined by Tanford (J. Am. Chem. Soc. 84:4240-4274(1962)) |
| 53 | Hydrophobicity_Welling | Total antigenicity, as defined by Welling (FEBS Lett. 188:215-218(1985)) |
| 54 | Hydrophobicity_Wolfenden | Total hydration potential at 25 °C, as defined by Wolfenden (Biochemistry 20:849-855(1981)) |
| 55 | Molecular_Weight | Molecular weight based on the sum of each amino acid molecular weight |
| 56 | Number_Of_Codons | Number of codons encoding each amino acid in the universal genetic code |
| 57 | Parallel_Beta_Strand | Parallel beta strand propensity, as defined by Lifson and Sander (Nature 282:109-111(1979)) |
| 58 | Percentage_Accessible_Res | Total molar fraction of accessible residues, as defined by Janin (Nature 277:491-492(1979)) |

| | | |
|---|---|---|
| 59 | Percentage_Buried_Res | Total molar fraction of buried residues, as defined by Janin (Nature 277:491-492(1979)) |
| 60 | Polarity_Grantham | Total polarity, as defined by Grantham (Science 185:862-864(1974)) |
| 61 | Polarity_Zimmerman | Total polarity, as defined by Zimmerman (J. Theor. Biol. 21:170-201(1968)) |
| 62 | Ratio_Hetero_End_Side | Total atomic weight ratio of hetero elements in end group to C in side chain (Science 185:862-864(1974)) |
| 63 | Recognition_Factors | Total recognition factor of each amino acid, as defined by Fraga (Can. J. Chem. 60:2606-2610(1982)) |
| 64 | Refractivity | Total refractivity index of each amino acid, as defined by Jones (J. Theor. Biol. 50:167-184(1975)) |
| 65 | Relative_Mutability | Total relative mutability (Ala=100), as defined by Dayhoff et al. (In "Atlas of Protein Sequence and Structure", Vol.5, Suppl.3 (1978)) |
| 66 | Theoretical pI* | Isoelectric point |
| 67 | Total_Beta_Strand | Total (antiparallel+parallel) beta strand propensity, as defined by Lifson and Sander (Nature 282:109-111(1979)) |
| 68 | Transmembrane_Tendency | Total transmembrane tendency, as defined by Zhao and London (Protein Sci. 15:1987-2001(2006)) |

**Table S2.** Summary of descriptor importance and model performance for a VH-based LogisticReg model trained on all descriptors (excluding charge at pH 6 and 7.4). For each descriptor, the following is provided; (i) cluster number assigned by hierarchical clustering of Spearman correlation coefficients as a way to group redundant descriptors, (ii) LogisticReg coefficients from the logistic regression model, (ii) absolute value of LogisticReg coefficients from the LogisticReg model, (iii) the decrease in model accuracy when the descriptor is permuted, indicating its importance (based on 10-fold CV on test data), (iv) the model accuracy when the specific descriptor is left out, indicating its unique contribution, and (v) the accuracy of the model using only the single descriptor. Descriptors are listed along with their respective cluster number and their importance metrics, highlighting their contribution to the model's performance and their individual predictive power.

| # | Descriptor | Cluster number | LogisticReg coeff. | LogisticReg abs(coeff.) | Permutation on test data [%-units decrease in accuracy] | Leave-one-feature-out accuracy [%] | Single-descriptor model accuracy [%] |
|---|---|---|---|---|---|---|---|
| 1 | Disorder_Propensity_DisProt | 7 | 0.51 | 0.51 | 2.9 | 70.0 | 50.2 |
| 2 | Disorder_Propensity_TOP_IDP | 12 | 0.46 | 0.46 | 2.5 | 69.8 | 48.8 |
| 3 | Theoretical pI | 22 | 0.45 | 0.45 | -3.5 | 68.6 | 65.2 |
| 4 | Aggrescan_av4 | 8 | 0.37 | 0.37 | -0.2 | 70.0 | 55.4 |
| 5 | Percentage_Accessible_Res | 6 | -0.36 | 0.36 | 1.4 | 70.1 | 52.0 |
| 6 | Zyggregator_profile_smoothed_pos | 14 | 0.35 | 0.35 | -1.7 | 69.5 | 57.1 |
| 7 | Hplc_Hfba_Retention | 8 | 0.33 | 0.33 | -4.3 | 70.1 | 57.3 |
| 8 | Hydrophobicity_Tanford | 8 | -0.31 | 0.31 | 0.3 | 70.0 | 53.1 |
| 9 | Hydrophobicity_Hplc_Ph_3_4_Cowan | 8 | -0.29 | 0.29 | 1.6 | 70.1 | 54.6 |
| 10 | Polarity_Zimmerman | 15 | -0.27 | 0.27 | 0.0 | 70.0 | 61.2 |
| 11 | Aggrescan_Nr_hotspots | 17 | -0.26 | 0.26 | 0.0 | 69.1 | 49.3 |
| 12 | Hydrophobicity_Hplc_Ph_7_5_Cowan | 8 | 0.25 | 0.25 | 0.8 | 70.0 | 57.6 |
| 13 | Beta_Sheet_Chou_Fasman | 10 | 0.24 | 0.24 | -0.6 | 70.0 | 54.8 |
| 14 | Hydrophobicity_Welling | 18 | -0.22 | 0.22 | -0.6 | 69.7 | 54.7 |
| 15 | Refractivity | 21 | -0.22 | 0.22 | 3.0 | 70.0 | 49.4 |
| 16 | Ratio_Hetero_End_Side | 19 | -0.21 | 0.21 | 2.1 | 70.0 | 53.1 |
| 17 | Average_Flexibility_BP | 2 | 0.21 | 0.21 | 1.1 | 70.0 | 54.6 |
| 18 | Beta_Turn_Chou_Fasman | 4 | -0.20 | 0.20 | 3.3 | 69.8 | 50.9 |
| 19 | Hydrophobicity_Hplc_Parker | 8 | -0.20 | 0.20 | 0.0 | 70.0 | 57.0 |
| 20 | Hydrophobicity_Roseman | 8 | 0.19 | 0.19 | 2.1 | 70.0 | 57.1 |
| 21 | Beta_Turn_Deleage_Roux | 5 | 0.19 | 0.19 | -0.6 | 70.0 | 55.3 |
| 22 | Aa_Composition_Swissprot | 9 | 0.19 | 0.19 | 0.6 | 70.0 | 52.9 |
| 23 | Hydrophobicity_Eisenberg | 8 | -0.19 | 0.19 | 0.8 | 70.0 | 53.7 |
| 24 | Number_Of_Codons | 9 | 0.19 | 0.19 | 2.7 | 70.0 | 56.8 |
| 25 | Bulkiness | 10 | 0.17 | 0.17 | -1.3 | 70.0 | 56.8 |
| 26 | Hydrophobicity_Sweet | 7 | 0.16 | 0.16 | 0.8 | 70.0 | 50.8 |
| 27 | Alpha_Helix_Levitt | 1 | 0.15 | 0.15 | -0.3 | 70.0 | 49.8 |
| 28 | Hydrophobicity_Janin | 8 | -0.13 | 0.13 | -0.2 | 70.0 | 54.1 |
| 29 | Beta_Turn_Levitt | 4 | -0.12 | 0.12 | 3.5 | 70.0 | 50.1 |
| 30 | Aa_Composition | 9 | 0.12 | 0.12 | 1.0 | 70.0 | 52.7 |
| 31 | Hydrophobicity_Hopp_Woods | 7 | -0.12 | 0.12 | 1.9 | 70.0 | 55.6 |
| 32 | Hydrophobicity_Fauchere | 8 | -0.12 | 0.12 | 1.9 | 70.0 | 54.7 |

| | | | | | | | |
|---|---|---|---|---|---|---|---|
| 33 | Hydrophobicity_Rao_Argos | 10 | -0.12 | 0.12 | 1.9 | 70.0 | 53.1 |
| 34 | Recognition_Factors | 2 | -0.11 | 0.11 | 3.2 | 70.0 | 53.2 |
| 35 | Alpha_Helix_Chou_Fasman | 1 | -0.11 | 0.11 | 2.1 | 70.0 | 50.4 |
| 36 | Hydrophobicity_Abraham_Leo | 8 | -0.11 | 0.11 | 2.7 | 70.0 | 52.4 |
| 37 | Aggrescan_av4_pos | 8 | 0.10 | 0.10 | 1.3 | 69.8 | 53.4 |
| 38 | Relative_Mutability | 20 | 0.09 | 0.09 | -1.3 | 70.0 | 50.9 |
| 39 | Hydrophobicity_Rf_Mobility | 10 | -0.09 | 0.09 | 2.7 | 70.0 | 54.4 |
| 40 | HPLC_Tfa_Retention | 8 | 0.08 | 0.08 | -0.5 | 70.0 | 53.2 |
| 41 | HPLC_Retention_Ph_2_1 | 7 | -0.07 | 0.07 | 3.2 | 70.0 | 54.9 |
| 42 | Hydrophobicity_Guy | 8 | -0.07 | 0.07 | 1.1 | 70.0 | 51.3 |
| 43 | Aa_Flexibility_VTR | 2 | 0.07 | 0.07 | -0.5 | 70.0 | 51.7 |
| 44 | Hydrophobicity_Manavalan | 10 | 0.06 | 0.06 | -1.0 | 70.0 | 52.8 |
| 45 | Hydrophobicity_Chothia | 10 | -0.06 | 0.06 | 2.4 | 70.0 | 55.1 |
| 46 | Hydrophobicity_Bull_Breese | 7 | 0.06 | 0.06 | 1.3 | 70.1 | 50.6 |
| 47 | Hydrophobicity_Miyazawa_Jernigan | 10 | 0.06 | 0.06 | -0.5 | 70.0 | 53.3 |
| 48 | Alpha_Helix_Deleage_Roux | 1 | -0.05 | 0.05 | 2.2 | 70.0 | 53.1 |
| 49 | Beta_Sheet_Levitt | 10 | -0.05 | 0.05 | 3.7 | 70.0 | 54.8 |
| 50 | Hydrophobicity_Black | 8 | -0.04 | 0.04 | 3.0 | 70.0 | 53.9 |
| 51 | Disorder_Propensity_FoldUnfold | 10 | -0.04 | 0.04 | 3.2 | 70.0 | 49.7 |
| 52 | Hydrophobicity_Kyte_Doolittle | 8 | -0.03 | 0.03 | 0.2 | 70.0 | 53.2 |
| 53 | Percentage_Buried_Res | 10 | -0.03 | 0.03 | 0.6 | 70.0 | 51.2 |
| 54 | Hydrophobicity_Hplc_Wilson | 13 | -0.03 | 0.03 | 3.0 | 70.0 | 47.6 |
| 55 | Polarity_Grantham | 23 | 0.02 | 0.02 | -1.4 | 70.0 | 52.0 |
| 56 | Total_Beta_Strand | 10 | -0.02 | 0.02 | 2.9 | 70.0 | 53.7 |
| 57 | Zyggregator_profile_smoothed | 14 | -0.02 | 0.02 | 1.7 | 70.1 | 57.3 |
| 58 | Parallel_Beta_Strand | 10 | -0.02 | 0.02 | 2.9 | 70.0 | 54.2 |
| 59 | Hydrophobicity_Rose | 10 | -0.02 | 0.02 | 2.7 | 70.0 | 51.0 |
| 60 | Antiparallel_Beta_Strand | 10 | -0.02 | 0.02 | 2.7 | 70.0 | 53.4 |
| 61 | Hydrophobicity_Wolfenden | 16 | 0.01 | 0.01 | 1.0 | 70.0 | 52.9 |
| 62 | Coil_Deleage_Roux | 3 | -0.01 | 0.01 | 1.6 | 70.0 | 52.8 |
| 63 | Hplc_Retention_Ph_7_4 | 7 | 0.01 | 0.01 | 0.6 | 70.0 | 56.5 |
| 64 | Beta_Sheet_Deleage_Roux | 10 | 0.01 | 0.01 | 0.3 | 70.0 | 55.1 |
| 65 | Transmembrane_Tendency | 8 | 0.00 | 0.00 | 0.0 | 70.0 | 54.9 |
| 66 | Avg_Area_Buried | 11 | 0.00 | 0.00 | 0.0 | 70.0 | 50.1 |

171
